# Transcription feedback dynamics in the wake of cytoplasmic degradation shutdown

**DOI:** 10.1101/2021.12.01.470637

**Authors:** Alon Chappleboim, Daphna Joseph-Strauss, Omer Gershon, Nir Friedman

## Abstract

In the last decade, multiple studies have shown that cells maintain a balance of mRNA production and degradation, but the mechanisms by which cells implement this balance remain unknown. Here, we monitored cells’ mRNA and nascent mRNA profiles immediately following an acute depletion of Xrn1 - the main 5’-3’ mRNA exonuclease - that was previously implicated in balancing mRNA levels. We captured the detailed dynamics of the cells’ adaptation to rapid degradation of Xrn1 and observed a significant accumulation of mRNA, followed by a delayed global reduction in nascent transcription and a gradual return to baseline mRNA levels. We present evidence that this transcriptional response is linked to cell cycle progression, and that it is not unique to Xrn1 depletion; rather, it is induced earlier when upstream factors in the 5’-3’ degradation pathway are perturbed. Our data suggest that the RNA feedback mechanism is cell-cycle-linked and monitors the accumulation of inputs to the 5’-3’ exonucleolytic pathway rather than its outputs.

## Introduction

Gene expression is a multistep process that starts at the nucleus, where structural (e.g., histones) and regulatory factors interact to facilitate transcription by the general transcription machinery and RNA polymerase II (PolII). During transcription, nascent mRNA molecules are capped at their 5’ end with a nucleolytic-resistant nucleotide (m^7^G), spliced, cleaved and polyadenylated. Protein-mRNA complexes are then exported from the nucleus to undergo translation in the cytoplasm by ribosomes. To allow for dynamic gene expression, most mRNA species are actively degraded by cells within a short time frame (minutes in yeast to hours in mammalian cells (Baudrimont et al., 2017; Yang et al., 2003)). mRNA degradation can be triggered by various quality control mechanisms (e.g. nonsense-mediated decay), but degradation is also thought to be coupled to translation of the mRNA by ribosomes (Hu, 2016; Huch and Nissan, 2014). Seminal work established the main degradation pathway in eukaryotes: mRNA is deadenylated by the Ccr4-Not complex, decapped by the Dcp1-Dcp2 complex, and degraded by the highly processive 5’-3’ exoribonuclease Xrn1 (Decker and Parker, 1993; Dunckley and Parker, 1999; Geisler and Coller, 2012; Hsu and Stevens, 1993; Muhlrad et al., 1994; Stevens, 1978). An important alternate route involves the exosome, which degrades mRNA from its 3’ end following deadenyation by Ccr4-Not complex. Additional less common endonucleolytic degradation pathways were also described (Decker and Parker, 1993; Łabno et al., 2016).

As presented above, this process is largely unidirectional, namely messages are generated in the nucleus, exported, translated and degraded with no information flow back to the nucleus. However, mounting evidence from the past decade suggests that transcription in the nucleus is coupled to degradation in the cytoplasm. This coupling was demonstrated along two main branches of evidence. The first is gene-specific regulation of transcript fate by nuclear signals, e.g. replacing the promoter of a gene can alter its transcript half-life or cytoplasmic localization (Bregman et al., 2011; Dori-Bachash et al., 2012; Slobodin et al., 2020; Trcek et al., 2011; Zid and O’Shea, 2014). In these cases, the functional implications are generally thought to be carried out by different mRNA-binding proteins that are exported with the transcript (Haimovich et al., 2013a).

Another line of evidence linking nuclear transcription and cytosolic degradation is of a more global nature (termed mRNA buffering or mRNA homeostasis). It was demonstrated that large scale perturbations to the degradation machinery as a whole are compensated by the transcription machinery (Haimovich et al., 2013b; He et al., 2003; Sun et al., 2013) and vice versa (Baptista et al., 2017; Dori-Bachash et al., 2012; Goler-Baron et al., 2008; Plaschka et al., 2015; Rodríguez-Molina et al., 2016; Shalem et al., 2008, 2011; Slobodin et al., 2020; Sun et al., 2012). In these cases, the underlying mechanisms for the observed global transcription-degradation coupling remain contested and speculative. Proposed mechanisms involve several main components, including Rpb4/7 (POLR2D/G in human) (Duek et al., 2018; Goler-Baron et al., 2008; Schulz et al., 2014), Pab1/Nab2 (Gilbertson et al., 2018; Kumar et al., 2011; Schmid and Jensen, 2018), the Ccr4-Not complex (Collart, 2016; Slobodin et al., 2020), Snf1 (AMPK) (Braun and Young, 2014; Young et al., 2012), and Xrn1 (Haimovich et al., 2013b; Sun et al., 2013).

Xrn1 is the main cytosolic 5’-3’ exonuclease in eukaryotic cells (Geisler and Coller, 2012; Heyer et al., 1995; Hsu and Stevens, 1993; Nagarajan et al., 2013; Parker, 2012). Xrn1 knockdown causes developmental and fertility defects in multicellular organisms (Jones et al., 2012; Newbury, 2004), and its knockout in yeast was shown to be detrimental to growth in certain stress conditions, to affect cell size and growth rate, and cause spindle-pole separation defects (Interthal et al., 1995; Jorgensen et al., 2002; Larimer and Stevens, 1990). In two important works published in 2013 Xrn1 was implicated in the coupling between degradation and transcription (Haimovich et al., 2013b; Sun et al., 2013). However, these studies arrived at opposite conclusions about the role of Xrn1. Haimovich and colleagues reported that Xrn1 knockout maintains global mRNA levels. They could explain the observed buffering by attributing to Xrn1 (and related RNA binding factors) a role as a transcriptional activator that acts directly on chromatin. Conversely, in a systemic screen of RNA processing factors, Sun and colleagues reported that Xrn1 knockout results in the most significant increase in total mRNA levels, which is the result of a significant decrease in degradation rates and a slight increase in global transcription rates. Mechanistically, they linked Xrn1 levels to a control over the transcript levels of a negative transcriptional regulator - Nrg1. Since then, other studies with Xrn1 knockout/knockdown exhibited various transcriptional effects (Begley et al., 2021; Blasco-Moreno et al., 2019; Chattopadhyay et al.; van Dijk et al., 2011; Fischer et al., 2020; García-Martínez et al., 2021a; Medina et al., 2014). Similarly, the Ccr4-Not complex which is crucial for mRNA deadenylation and degradation was also implicated in transcription regulation (Collart, 2016; Collart and Struhl, 1994; Gupta et al., 2016; Slobodin et al., 2020). Interestingly, in the systemic screen by Sun et al. some components of the Ccr4-Not complex incur pronounced deviations from the wildtype strain (Sun et al., 2013).

To summarize, there is ample evidence that mRNA buffering takes place in perturbed yeast cells, and probably in higher eukaryotes, but there is little understanding of the mechanisms underlying various contradicting observations. We reasoned that if a feedback process takes place, the dynamics of the process can shed light on its mechanistic details. However we found little information about the dynamics of the process, as most studies were performed in knockout strains at steady-state conditions. In several notable exceptions the time scale of the feedback was determined to be in the order of several minutes(Baptista et al., 2017) (baptista) and upto an hour (Plaschka et al., 2015; Sun et al., 2013), but these were observed in different settings, and their generality is unclear.

Several of the works mentioned here applied comparative Dynamic Transcriptome Analysis (cDTA (Sun et al., 2012)) to samples, which measures mRNA levels and monitors nascent transcription by pulse labeling RNA with uracil analogs. We adapted the cDTA protocol to a high-throughput, quantitative, and sequencing-based version that we termed cDTA-seq. The cDTA-seq technique allowed us to monitor Xrn1 depletion from cells in high resolution, and observe the dynamics of mRNA accumulation and cells’ adaptation to this perturbation. Utilizing metabolic labeling data we identify a delayed global reduction in nascent transcription, which results in the return to wildtype mRNA levels after several hours. By repeating this dynamic experiment in G1-arrested cells, and by analyzing cell-cycle transcripts in unsynchronized populations, we propose that the delayed transcriptional response is linked to the cell cycle. We further expanded our data to multiple other RNA processing factors and found that the adaptive response was not unique to Xrn1, and it also occurred upon depletion of Dcp2 and Not1. Interestingly, we find that the transcriptional response initiates earlier when upstream components in the 5’-3’ pathway are perturbed, suggesting that the trigger for the transcriptional response is sensed upstream of the degradation pathway. These results provide a rich resource for studying the cellular response to perturbations in the general mRNA degradation machinery, and our analysis provides insights to basic properties of the mechanism underpinning the mRNA buffering phenomenon.

## Results

### High-throughput, quantitative dynamic transcriptome analysis by sequencing (cDTA-seq)

To study the dynamic equilibrium of mRNA in cells a dynamic measurement is needed. We aimed to assess detailed time course absolute mRNA levels and transcription rates in multiple strains, resulting in hundreds of samples. Therefore, we developed a protocol (Figure 1A) inspired by Sun and colleagues (Sun et al., 2012), in which we spike-in a known amount of cells from a close species for mRNA quantification, combined with a brief pulse with 4-thiouracil (4tU) to quantify recently transcribed molecules. Following recent works, we alkylate the 4tU nucleotides with iodoacetamide (Herzog et al., 2017; Voichek et al., 2016), which results in their subsequent reverse-transcription to G instead of A (T→C on the sense strand). The conversion allows for detection of 4tU at the same time as quantifying mRNA levels with RNA-seq. Thus, we avoid additional biochemical separation and unknown losses in each sample, which allows for accurate estimation of the fraction of nascent molecules (Figure 1B). We incorporated these steps into an RNA extraction and 3’ mRNA sequencing protocol we had recently developed that was specifically aimed at high throughput measurements (Klein-Brill et al., 2019). The final protocol (Figures 1A-B) allows a single person to quantify the entire transcriptome, including nascent mRNA, from 192 samples in a single day (methods).

**Figure 1.**
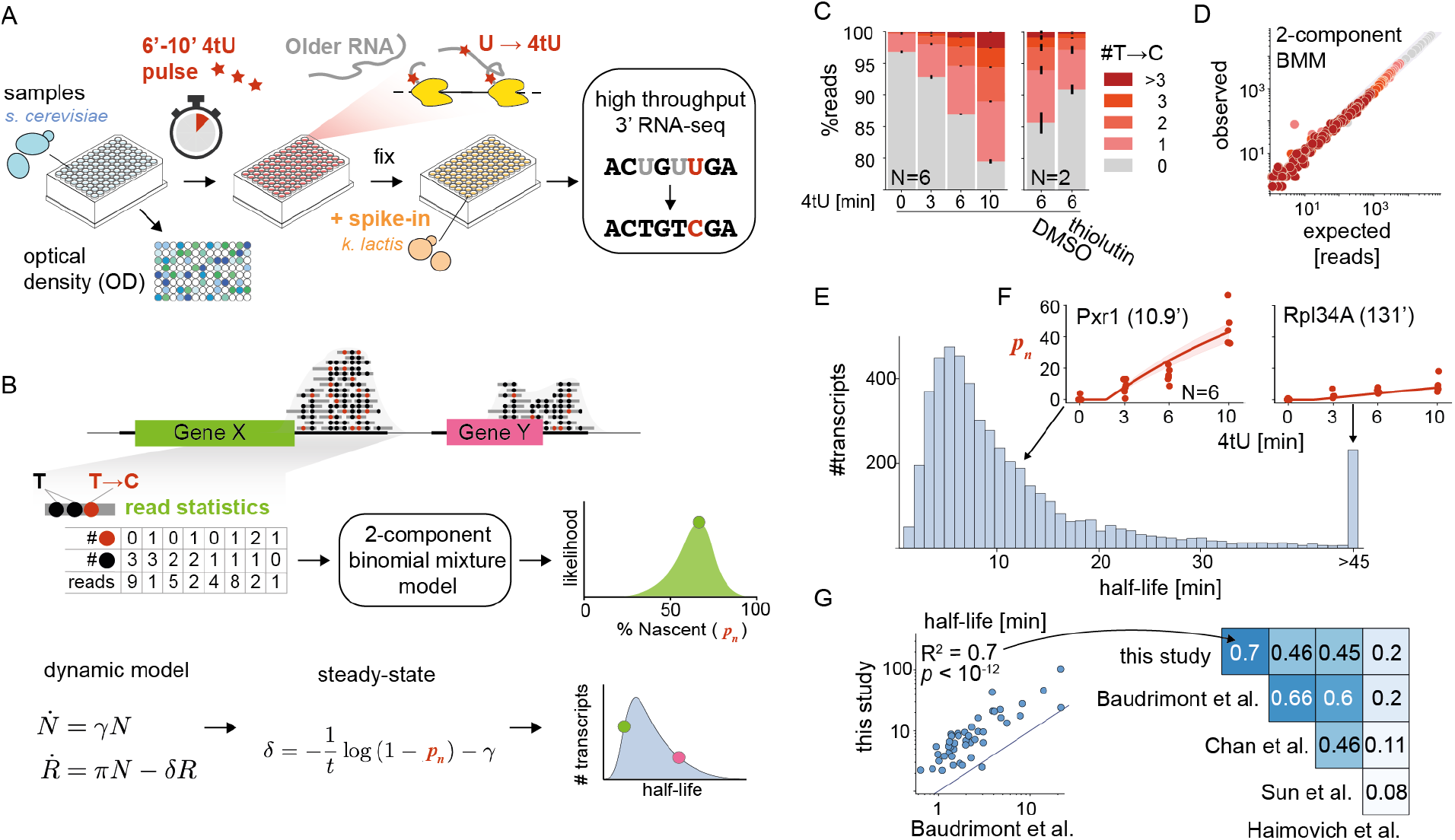
cDTA-seq and genome-wide transcript half-life estimation. **A) cDTA-seq protocol outline.** 4tU is added to dozens of quantified samples to label nascent RNA. Cells are then immediately fixed with a pre-fixed constant amount of spike-in cells. RNA is extracted, 4tU is alkylated and RNA-seq libraries are prepared, resulting in T→C conversions where 4tU was incorporated. The entire process is performed in a 96 sample format. **B) Transcript-level analysis outline.** Read conversion statistics per gene are fitted with a binomial mixture model to estimate the percent of nascent molecules (*p_n_*). Assuming a first-order kinetic model (N - number of cells, R - number of mRNA molecules, (*γ*, *π*, *δ*) are growth, production, and degradation rates respectively), and assuming steady-state, *p_n_* can be translated to transcript half-life given the known labeling period (*t*). See methods for more details. **C) 4tU conversion is effective, reproducible and measures transcription.** Percent of reads (y-axis) along a 4tU time course (x-axis) with a different number of observed T→C conversions (legend, N = 6). Samples exposed to vehicle (DMSO) or the transcription inhibitor thiolutin for 15 minutes (and labeled with 4tU for 6 minutes, N = 2). Note that the y-axis begins at 75%, i.e. most reads have no conversions. **D) Binomial Mixture Model (BMM) fits the data.** Read conversion statistics are fitted with a 2-component BMM. Each dot represents the number of reads with a certain number of observed Ts and T→C conversions (color as in E). x-axis is the expected number of reads for each (T,T→C) pair assuming the model and the observed #T distribution in the data, y-axis is the observed number of reads in each (T,T→C) combination (grey highlight is x = y). Additional components do not improve the likelihood of the data (Figure S1D). **E) Half-life distribution for all yeast transcripts.** Assuming steady-state, global and transcript specific parameters are iteratively fitted, resulting in the half-life of each transcript. The median of the distribution is 8.2 minutes, transcripts with a half-life of 45’ or longer are counted in the rightmost bin. **F) Examples of estimated *p_n_* along the time course.** The individually estimated *p_n_* per time point and replicate (N = 6) for two transcripts (Pxr1 and Rpl34A) is shown as red dots along the time course. The data is fitted with a single parameter per gene (degradation rate) resulting in an estimate of 10.9’ half-life for Pxr1 and a 131’ half-life for Rpl34A. Using these estimates, the expected *p_n_* along the time course is plotted as a red line with 95% CI as a red shaded area. **G) Half-life estimates correlate with various studies.** Half-lives from this and four other published studies are compared to each other and the linear explanatory value (R^2^) is denoted in the upper diagonal matrix. The scatter depicts a specific example of the comparison between this study and the estimates from Baudrimont et al where estimates were not obtained by metabolic labeling.

We verified our ability to quantify sample mRNA levels by spike-in titration (Figure S1A, R^2^ = 0.99, *p* < 10^-13^). To avoid manual cell counting on hundreds of samples, we validated that optical density (OD) was a good proxy for the number of cells in various genotypes and conditions used in this study (as represented by DNA content, see methods, Figure S1B, R^2^ = 0.89, *p* < 10^-47^).

Next, we performed a 4tU labeling time course experiment and observed a significant increase only in T→C conversions (Figures 1C, S1C). To verify that we can detect changes in transcription rates, we also performed this analysis after transcription inhibition with thiolutin for 15’ and pulse-labeled the samples with 4tU to observe a >50% reduction in labeled molecules (Figure 1C). As previously observed (Jürges et al., 2018), individual reads exhibit multiple conversion events (Figure 1C) bolstering confidence that they arise from newly transcribed molecules. We used this observation to estimate the percent of molecules that were transcribed during the labeling period (*p_n_*) by fitting a probabilistic model (binomial mixture model, methods) to the data (Figure 1B (Jürges et al., 2018)). We verified that the model fitted the observed data well (Figures 1D, S1D), and that the data is best explained by an increase in the proportion of nascent molecules (*p_n_*), rather than e.g. changes to incorporation efficiency or other artifacts (Figure S1E).

Assuming a steady-state and a first order model for mRNA (Figures 1B, S1F-H), a half-life can be calculated per transcript (Figures 1E-F, S1I-J). We compared our estimates with four published works that measured or estimated transcript half-lives using various methods (Figure 1G) (Baudrimont et al., 2017; Chan et al., 2018; Haimovich et al., 2013b; Sun et al., 2013). Our technique is clearly correlated to other studies, but there are global discrepancies between half-life estimates that were previously noted (Figure S1K) (Harigaya and Parker, 2016; Sun et al., 2012; Wada and Becskei, 2017), that are potentially explainable by technical discrepancies (e.g. maturation and polyadenylation time can cause an offset between techniques, Figures S1L).

We conclude that cDTA-seq can be used for relative mRNA and half-life estimation per transcript from a single measurement, with the caveat that the absolute half-life numbers from any technique should be treated with care. Thus, in each experimental batch we include a wildtype sample, which is used as a control for physiological anomalies and technical discrepancies between experiments.

### Turnover rates are slowed in the absence of Xrn1, but Global mRNA levels are maintained

Previous studies found Xrn1 knockout to cause a global increase in mRNA levels per cell (Sun et al., 2013), or conversely, an unchanged level of mRNA (Fischer et al., 2020; Haimovich et al., 2013b). To address the question of absolute mRNA levels, we applied cDTA-seq to quantify the changes in mRNA levels in the absence of Xrn1 and the underlying mRNA dynamics. Given that Xrn1 is a major mRNA degradation factor, a knockout of Xrn1 is expected to cause an increase in total mRNA levels. However, using our protocol we do not observe any difference in global mRNA levels when comparing fresh Δxrn1 strains to wt cells (Figure 2A). Further, despite significant changes to ~400 transcripts (Figure S2A), the overall distribution of mRNA was unchanged (Figure 2B). We then turned to examine the changes to transcription and degradation rates in the absence of Xrn1. Contradicting reports in the literature argue that transcription slightly increases (Sun et al., 2013)) upon Xrn1 knockout, or is markedly reduced (Haimovich et al., 2013b). In our data (two biological replicates in two fresh knockouts) we find a significant reduction in the fraction of nascent molecules from 22.8% to 14% (Figure 2A, t-test *p* < 0.004). Using our data, we were able to estimate the degradation and transcription rates for ~4800 transcripts (>70% of genes). Unlike mRNA levels, we do observe a clear reduction in degradation rates (median decrease of 40%, with 746 transcripts becoming completely stable, Figure 2C). A corresponding reduction in transcription rates (median decrease of 70%, Figure S2B) can be inferred from the mRNA levels and degradation rate estimates. When we examined the relationship between changes in production and degradation rates per transcript we found strong agreement, consistent with buffering (*r* 0.87, *p* < 10^-300^, Figures 2D, S2C).

**Figure 2.**
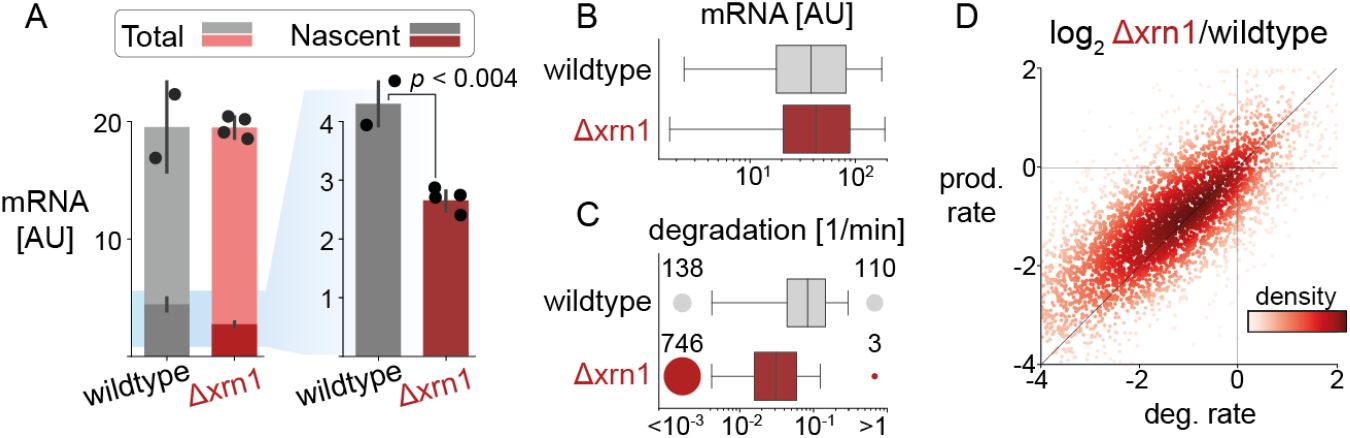
Xrn1 knockout causes a genome-wide decrease in degradation and transcription rates but maintains global mRNA levels. **A) Global mRNA levels are maintained while nascent fraction decreases significantly.** The amount of total mRNA (y-axis) in wildtype and Δxrn1 is the same (cumulative bars), while the fraction of nascent molecules decreases significantly (t-test *p* < 0.004, dark bars). **B) mRNA distribution is relatively unchanged between Xrn1 and wildtype.** Transcript abundance distribution in wildtype (grey) and Δxrn1 (red). Boxes throughout the manuscript mark the interquartile range (IQR) with whiskers at 1.5 × IQR. **C) Xrn1 knockout causes a transcriptome-wide decrease in degradation rates.** Transcript degradation rate distributions. Circles (and numbers) to the left and right of boxes correspond to the number of transcripts that are too stable (half-life > 3 hours, left) or too volatile (half-life < 1 min, right) to be estimated confidently. **D) Changes to degradation and production rates are correlated.** Production rates are inferred from mRNA levels and estimated degradation rates (methods). Log2 fold changes between Δxrn1 and wildtype in production rate (x-axis) and degradation rate (y-axis) per transcript (dots, colored by their local density). Pearson *r* = 0.87, *p* < 10^-300^.

To better understand the global changes to degradation and transcription rates, we looked for functional signatures in ours and in published Δxrn1 transcriptome data (Celik et al., 2017; Kemmeren et al., 2014; Sun et al., 2013). In these studies there is not data pertaining to absolute mRNA levels, but strikingly, in all three published datasets, Δxrn1 exerted the most pronounced change to the mRNA profile relative to wildtype (Figure S2D), and these profiles are correlated (Figure S2E). However, we could not identify robust common targets or consistent functional enrichments in the sets of up- and down-regulated genes in the knockout studies (Figure S2F). Similarly, we found that in our data virtually only ribosomal protein genes and ribosome biogenesis genes were exceptional, pointing to indirect growth effects (Figure S2G). Having excluded functional explanations we tried various gene/transcript features and sequence information to explain observed changes in mRNA levels or degradation and transcription rates. However, our models only explained a small fraction of the observed variance (Figures S2H-J, supplementary note).

We conclude that cells maintain their global mRNA levels in the absence of Xrn1 by reducing global transcription rates, but the specific details underpinning the homeostasis are obscure in the knockout strain.

### Conditional Xrn1 depletion reveals a global but transient increase in mRNA levels

We reasoned that if Xrn1 is a major exonuclease of mRNA (Geisler and Coller, 2012; Sun et al., 2013), it must exert an effect that is somehow buffered by cells. To test this hypothesis we set out to repeat the cDTA-seq experiment in an Xrn1 conditional knockdown strain. We generated strains in which Xrn1 is tagged with an Auxin Inducible Degron (AID) (Morawska and Ulrich, 2013; Nishimura et al., 2009) and validated that Xrn1 is depleted rapidly from cells (Figure 3A). To monitor the effects of Xrn1 depletion, we grew cells to exponential growth phase and performed a detailed 4-hour time-course experiment after Xrn1 depletion (Figure 3B). At the end of the time course, samples were simultaneously labeled with a short 4tU pulse and subjected to cDTA-seq to monitor their mRNA levels and nascent transcription at each time point following Xrn1 depletion.

**Figure 3.**
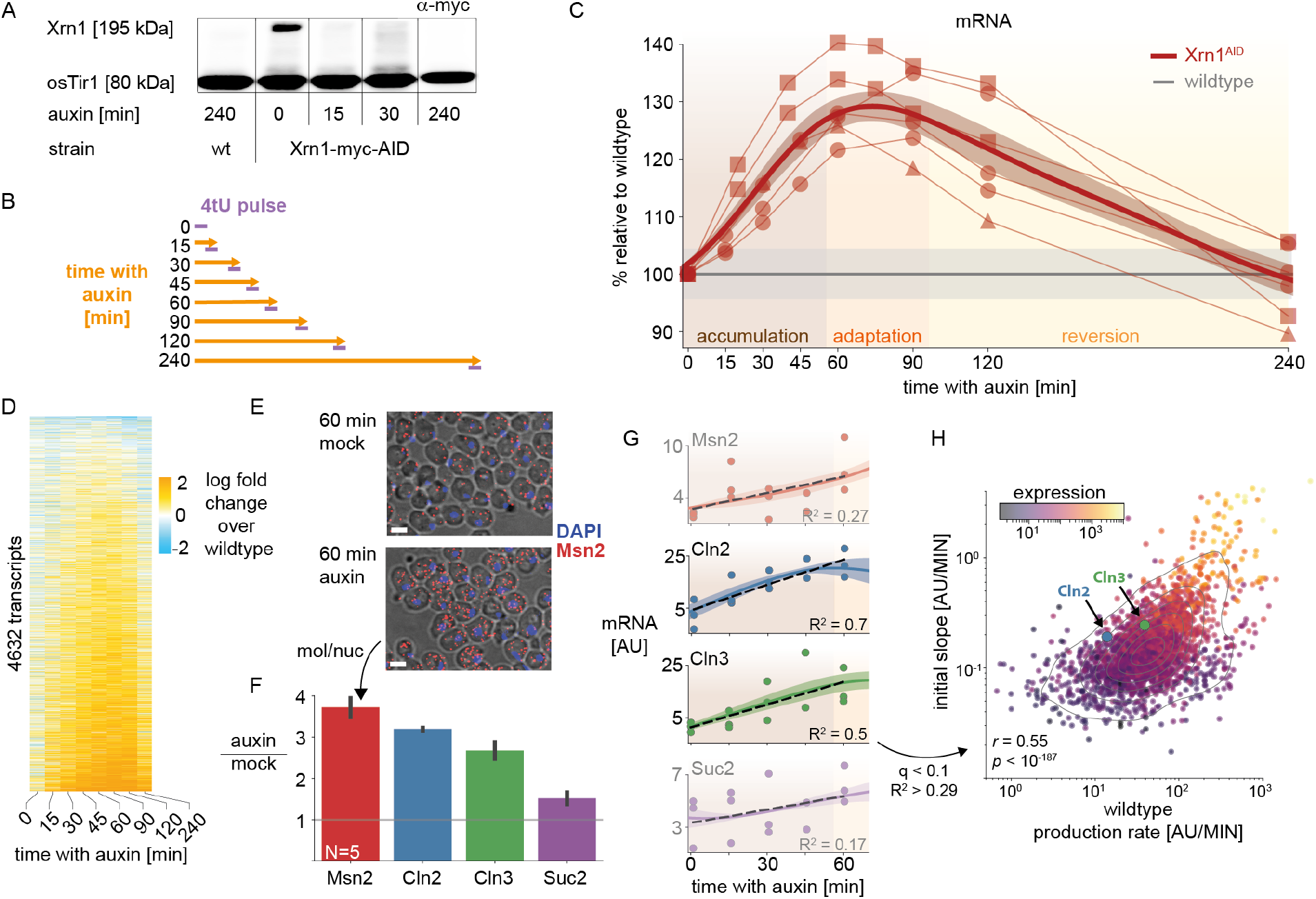
Xrn1 acute depletion causes a transient increase in mRNA levels. **A) Auxin inducible degradation (AID) of Xrn1 is rapid and stable.** Western blot (anti-myc) for Xrn1 tagged with an auxin inducible degron (AID) and a myc-tag. Shown are the isogenic untagged strain (left), and a time course that demonstrates virtually no Xrn1 protein within 15 minutes. osTir1 is also tagged with Myc in these strains and is used as a loading control. **B) AID/cDTA-seq experimental scheme.** Cells are grown to mid-log phase, and split. At each indicated time-point auxin is added to one culture, and after 4 hours (240 minutes) all samples are subjected to a short 4tU pulse simultaneously and harvested for cDTA-seq. **C) Accumulation of mRNA immediately following Xrn1 depletion.** mRNA counts (scaled by spike-in reads) from time course experiment (B, x-axis) in Xrn1^AID^ cells (red) are scaled to initial values, and to the corresponding wildtype measurement (y-axis) in three different experiments (markers). Grey line indicates the wildtype trajectory under the same transformation with standard deviation as a shaded area. Thick red line represents the average of the smoothed interpolations of each separate trajectory. We label the three stages of the response for convenience (accumulation → adaptation → reversion). **D) mRNA accumulation and reversion observed genome-wide.** mRNA per transcript (y-axis) was normalized with the corresponding measurement from the wildtype time course (time along x-axis), and the log fold change is color-coded. Transcripts were filtered to not have any missing values along the trajectory (N = 4632, ~70%), and were sorted by their average log fold change between 15 and 120 minutes. **E) single molecule FISH validation.** Composite micrographs (from a confocal Z-stack image), showing DIC image of cells with a max-projection of DAPI stain in blue and fluorescent probes for Msn2 in red (1 μm scale bar). Individual molecules and nuclei are clearly discernible and are counted. Xrn1^AID^ cells were treated with auxin or mock (DMSO) for 60 minutes, fixed, stained and imaged. A clear increase in molecule counts is observed. **F) single molecule FISH quantification.** In each field (N = 5), the number of observed molecules is divided by the number of observed nuclei to estimate the mRNA content of each cell in four different probes (x-axis). The y-axis denotes the ratio of the mRNA content in auxin vs. mock treatment. **G) scaled RNA trajectories, and rate of accumulation.** Each subplot shows the (spike-in scaled) mRNA counts (y-axis, points) from three replicates in the same experiment along the time course following Xrn1 depletion (x-axis). Line is the mean (+SEM) over interpolations of each separate repeat (N = 3). Selected transcripts correspond to smFISH probes (E-F). Dashed black line indicates the linear fit for the accumulation phase of the response (fit R^2^ noted). Only fits within the 10% FDR threshold (R^2^ > 0.29) are plotted in (H), Msn2 and Suc2 are shaded and do not appear in (H) as they are below this threshold. **H) mRNA accumulation correlates to transcript production rate.** Comparing slopes from the FDR-selected linear fits (y-axis, (H)), to the production rate estimated from the wildtype sample (Figure 2D), 30% of the observed variability (R^2^) can be attributed to the production rate (*p* < 10^-187^). Note that there is a strong correlation between the two measures and the overall expression (color-coded, log scale), see text and Figures S3E-F.

Multiple replicates reveal that mRNA levels increase significantly following Xrn1 depletion (20%-40% across multiple experiments and repeats; Figure 3C). However, after about 70 minutes from the time of auxin addition, the accumulation trajectory inverts, and we observe a decrease in global mRNA levels, resulting in a return to basal mRNA levels (90%-105%) within four hours. Importantly, the mRNA profile also converges to the Xrn1 knockout mRNA profile (Figure S3A), validating the effectiveness of Xrn1 depletion. We call the initial time period (0~55 min) the *accumulation* phase, the subsequent time (55~95 min) the *adaptation* phase, and during the final *reversion* phase cells settle back to their initial mRNA levels (95 min and onwards).

An examination of the mRNA profile along the time course reveals that it is not the result of an increase in specific highly-expressed transcripts, but that virtually all transcript levels are transiently increased (Figure 3D). We validated the increase in mRNA levels following 60 minutes of mock or auxin treatment by single molecule RNA-FISH with probes targeting four different transcripts, and recapitulated the results from our spike-in-normalized mRNA-seq counts (Figures 3E-F,S3B). Notably, in a different time course FISH experiment we observed that the reduction in smFISH signal is slower than the observed reduction in the cDTA-seq signal, and we have evidence to suggest that this is a result of the previously reported accumulation of non-polyadenylated transcripts in Xrn1-depleted cells, which are not captured in the cDTA-seq protocol (Figures S3C-D) (Hsu and Stevens, 1993).

These results highlight the ubiquitous nature of degradation by Xrn1 as evident by the immediate increase in the vast majority of transcripts, and reveal the dynamics of the cellular response to aberrant mRNA accumulation.

### mRNA accumulation correlates to transcription rates

While the response to Xrn1 depletion seems ubiquitous there are significant and reproducible differences between the response rate of different transcripts (Figures 3E-G). Given the direct nature of the perturbation, we wanted to test whether the observed changes to mRNA levels are consistent with a first order model for mRNA, whereby the immediate change in mRNA following a decrease in degradation is a function of individual transcript production rate (supplementary note). To test these predictions, we fit each transcript with a linear model for the change in mRNA during the accumulation phase of the response (Figure 3G). As predicted, the change in many transcripts is consistent with a linear increase (Figure S3E, 10% FDR, R^2^ > 0.29, 2,383/5,164 transcripts), and where we cannot reject the null hypothesis of constant levels, it is mostly due to sampling noise (low expression levels, see Msn2 and Suc2 in Figure 3G). We tested the correlation of the fitted slopes to the pre-perturbation transcription and degradation rates and found that the strongest correlation is to the production rate, as expected by a first order model (Pearson *r* = 0.55, *p* < 10^-187^, Figures 3I, S3F). To account for the potentially confounding effect of expression levels, which are correlated to both production rate and the slope (Figure 3H), we also calculated the conditioned correlation, which remained significant (partial Pearson *r* = 0.28, *p* < 10^-45^, Figure S3E). Furthermore, this dependency is significantly accentuated in transcripts with longer half-life (Figure S3G), consistent with masking effects of residual degradation following Xrn1 depletion.

Therefore, during the accumulation phase, mRNA increase is consistent with a scenario where degradation is abruptly reduced, while transcription remains largely unchanged which affects the rate of initial mRNA accumulation.

### Metabolic labeling through Xrn1 depletion uncovers a delayed global transcriptional adaptation

During the adaptation phase mRNA levels stop increasing and eventually revert back to WT levels. How does this occur within a few hours? Since mRNA level is at a balance of transcription, degradation, and dilution by growth, there are multiple possible explanations.

Cells can adapt to reduced Xrn1-dependent degradation rates by activating alternative mRNA degradation mechanisms, increasing their dilution rate, which requires faster cell division and growth, decreasing transcription rates, or respond in some combination of these mechanisms.

We first tested the hypothesis that cells adapt by increasing their division rate or volume. We monitored various physiological aspects of cellular growth following Xrn1 depletion and found no significant inflection points in optical density, cell size, cell counts, colony forming units, or cell-cycle fractions within two hours of auxin addition (Figure S4A-E). Notably, some of these measures do eventually change, as previously reported for the knockout strain (e.g. increase in cell size, Figure S4B). We also measured the growth rate in the absence of Xrn1 (knockouts and AID strains) and found a slower growth rate by 20-30% (Figures S4F-G). We therefore exclude the possibility of volume or growth increase as possible explanations for the observed adaptation in mRNA levels within two hours.

Another possibility is a compensatory increase in Xrn1-independent degradation. Xrn1 is the main 5’-3’ RNase in the cytoplasm, so it is possible that the 3’-5’ degradation branch is compensating for its absence. To test this hypothesis, we AID-tagged two components of the SKI complex (Ski2, Ski8) that were shown to be required for 3’-5’ degradation by the exosome (Anderson and Parker, 1998) in addition to the Xrn1 AID tag. While these double-AID strains exhibited significantly slower growth when exposed to auxin (Figures S4F-G), their immediate mRNA response to depletion was virtually identical to the Xrn1-AID strain (Figure S4H), suggesting that the 3’-5’ degradation branch, as mediated by SKI complex, does not take an active part in the observed reduction in mRNA levels.

To test the remaining possibility of transcriptional reduction, we turned to the nascent transcription data we collected. During the adaptation phase (55’-95’ following Xrn1 depletion) we observed a concerted and significant reduction in nascent transcripts (Figures 4A-B). Indeed, even genes that were induced immediately following Xrn1 depletion show a significant decrease at this point (Figure 4C), suggesting a global repression of transcription, which we term the *transcription adaptation response*.

**Figure 4.**
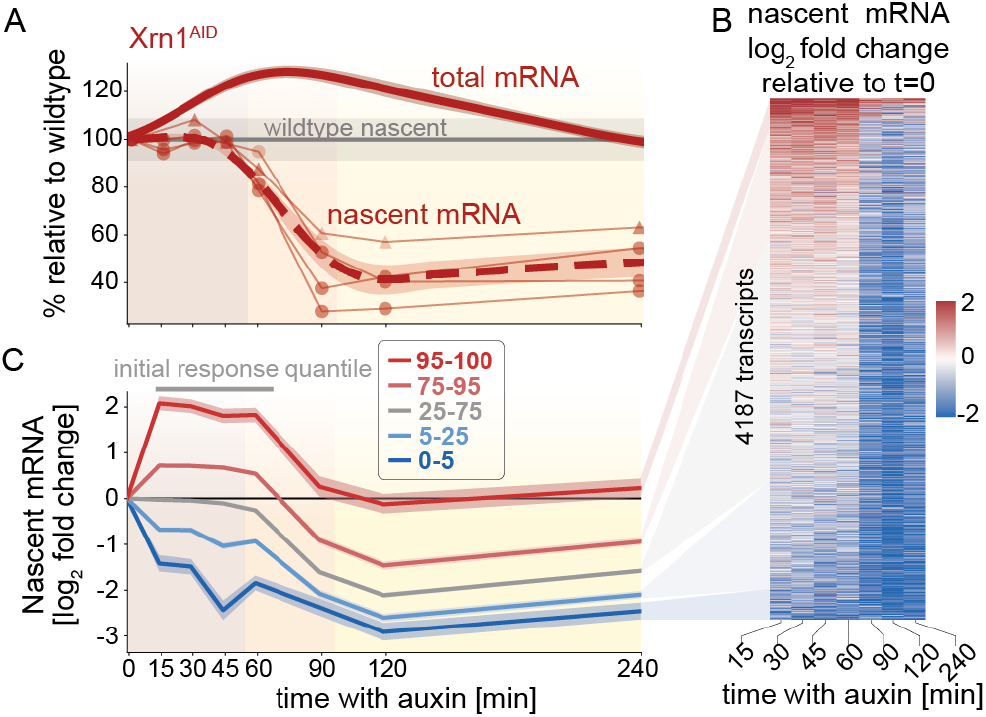
The transcription adaptation response to Xrn1 depletion. **A) Nascent mRNA is reduced by ~50% after ~60 minutes.** Global nascent counts (thin lines, N = 4) relative to t=0 from two experiments (markers) are plotted as a function of time since auxin addition (x-axis, shared with (C)). Dashed red line represents the average of the smoothed interpolations of each separate trajectory (N = 4). Grey line indicates the wildtype trajectory under the same transformation with standard deviation as a shaded area. Solid red line is the change to total mRNA levels (same as in Figure 3C). **B) Nascent reduction is abrupt and observed genome-wide.** Color-coded nascent mRNA log fold change (color scale) relative to t=0 per transcript (y-axis) along the Xrn1 depletion time course (x-axis, excluding t=0). Transcripts were filtered to not have any missing values along the trajectory (N = 4187, ~63%), and were sorted by their average log fold change between 15 and 60 minutes. **C) Transcription reduction is evident even in genes that were initially upregulated.** Transcripts were grouped by their average initial change (15 < t < 60, grey bar above plot) into percentile groups (legend), and the average log fold change trajectory of each group (y-axis) was plotted as a function of time since auxin addition (x-axis).

### The transcription adaptation response is linked to the cell-cycle

Even though the population of cells we studied was unsynchronized, the time scale of 60-90 minutes was in the order of the yeast cell cycle (~90’). To explore a potential role for the cell cycle in the transcription adaptation response, we performed cDTA-seq on samples from an Xrn1 depletion time-course in cycling and G1-arrested cultures (Figure 5A). When we compared the response to Xrn1 depletion in arrested/cycling cells, the accumulation of mRNA is evident and highly significant in both cycling and arrested cells (median increase of 46.3% and 22.9% increase respectively after 60 minutes, Fig 5B). Furthermore, changes to mRNA through the time-course are similar between cycling and arrested cells (Figures 5C, S5A), indicating that the depletion of Xrn1 induces the same immediate response even though arrested cells begin at a different basal state (Figures S5B-C). However, when we examined the transcription adaptation response, we observed only a minimal reduction in nascent transcription in arrested cells compared to cycling cells (Figures 5B-C). To exclude a possible concern that we are unable to reliably measure a decrease in nascent transcription relative to the initial arrested state, we depleted Med14 - an essential component of the mediator (Warfield et al., 2017) - which causes a significant reduction in nascent transcription in arrested cells (Figure S5D).

**Figure 5.**
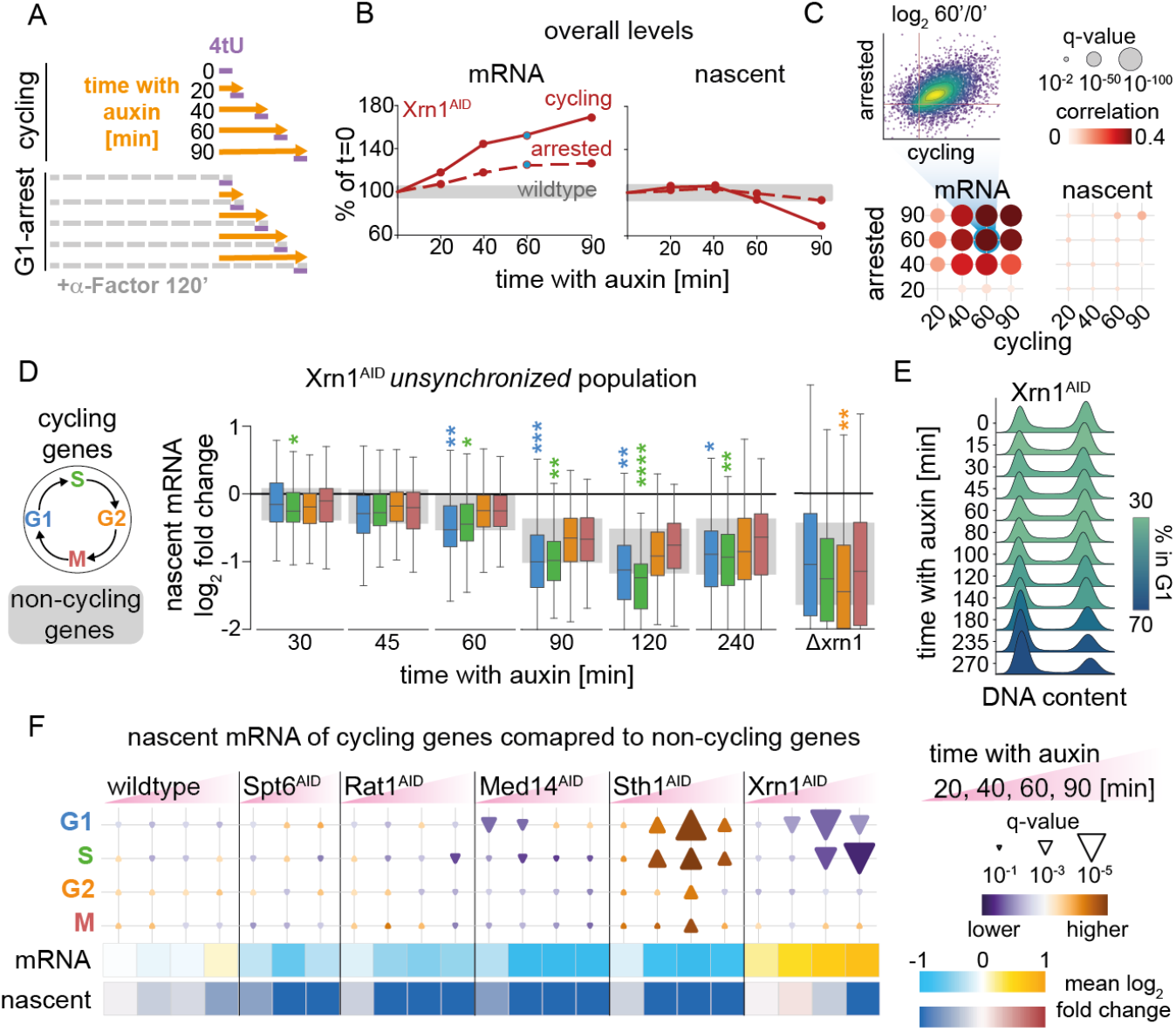
The transcription adaptation response is linked to the cell-cycle. **A) Cell-cycle arrest AID/cDTA-seq experimental scheme.** Cells are grown to mid-log phase and split to two cultures - with or without **α**-factor (a pheromone that arrests cells in G1). Cultures are then further split and auxin is added at indicated time points. All samples are subjected to a short 4tU pulse simultaneously and harvested for cDTA-seq (similar to Figure 3B). **B) Global changes to total and nascent mRNA following Xrn1 depletion in cycling and arrested cells.** Average change relative to t=0 (y-axis) as a function of time (since auxin addition, x-axis), in cycling (solid line) and arrested (dashed) cells. Changes to total mRNA on the left (60’ comparison shown in (C)), and changes to nascent mRNA on the right. Grey rectangle denotes the maximal deviation observed in the corresponding average (+SEM) calculated in the wildtype strain. **C) Correlations between cycling and arrested RNA profiles following Xrn1 depletion.** Dot plots denote correlations between different timepoints along the two time course experiments (x-axis: cycling, y-axis: arrested). Color proportional to Pearson *r*, dot size proportional to q-value (see legends on top-right). A comparison of the 60’ time point is shown as a scatter; each dot is a different transcript (colored by local density). Axes are the log2 fold change relative to t=0, which range between a 2-fold reduction and 4-fold increase both axes, Pearson *r* = 0.42, *q* < 10^-300^). **D) Cell-cycle signature following Xrn1 depletion in an unsynchronized population.** Using data from the experiment shown in Figures 3 and 4, we compare the distributions of changes to nascent transcription (relative to t=0, i.e. no auxin, x-axis) of each set of cycling genes (colors, legend) (Santos et al., 2015) to the distribution of non-cycling transcripts (shaded grey background is the IQR). Different time points in this dataset are organized from left to right (x-axis labels). Significant deviations (kolmogorov-smirnov q-values) are marked with 1/2/3/4 colored stars, if their q-value is smaller than 10^-2^, 10^-5^, 10^-10^, and 10^-15^ respectively. On the right - the same analysis applied to the Δxrn1 data shown in Figure 2. **E) FACS analysis of DNA-stained unsynchronized Xrn1^AID^ cells exposed to auxin.** DNA staining does not show any gross cell-cycle differences in the first 140 minutes. Only after 3 hours a shift to G1 is observed. Events histograms (y-axis) per 525nm filtered signal (x-axis, DNA was stained with SYBR green) are ordered along the time course following auxin addition (top to bottom), and are colored according to the percent of events associated with G1 phase DNA content (methods). **F) The cell cycle signature is unique to Xrn1 depletion.** We repeated the time course experiment on multiple different AID-tagged proteins. Differences in log fold change to nascent mRNA between cycling gene sets (row) and non cycling genes denoted as colored triangles (purple, down - lower than cycling genes, orange, up - higher, size proportional to kolmogorov-smirnov q-value). Each triangle denotes the difference along a specific depletion time point (columns, x-axis time since auxin addition, same as in (A-C)). Bottom panels denote the average log fold change to mRNA and nascent mRNA in the same samples.

While the differences between arrested and cycling cells were substantial, we were limited in our ability to probe the response to Xrn1 depletion for long durations in arrested cells due to their disrupted physiology and eventual escape from induced G1-arrest. We reasoned that if there is a relationship between cell-cycle and the transcription adaptation response it might be observable in data obtained from unsynchronized populations. To perform this analysis we used a classification of ~650 cycling genes to the four main stages of the cell cycle (Santos et al., 2015). We compared the mRNA distributions of cycling to non-cycling genes between wildtype and Δxrn1 cells and found no significant differences (Figure S5E). This suggested that Δxrn1 cells are able to adapt their cell-cycle transcripts to wildtype levels.

Next, we reasoned that if cells adapt to the absence of Xrn1, and this adaptation is linked to the cell cycle, there might be a transient cell-cycle signal following Xrn1 depletion that dissipates as cells adapt. To test this prediction, we used our measurements following Xrn1 depletion in an unsynchronized population, and specifically the changes to nascent transcription. While the cell-cycle gene sets start off as all other genes, we observed a faster and more pronounced transcription reduction in cycling G1 and S genes compared to other cell cycle groups or the rest of the transcriptome (kolmogorov-smirnov *q* 10^-2^-10^-18^, Figure 5D). As expected from an adaptive response, this signature dissipates as the cells arrive at their new steady-state. Following this observation, we monitored the cell-cycle distribution of an unsynchronized population following Xrn1 depletion (Figure 5E), and observed gross changes in DNA content distribution only after three hours. Thus, the earlier and subtle reduction to nascent transcription of G1 and S genes cannot be explained by changes to the distribution of cells along the cell cycle.

To verify that this cell-cycle signature is not an artifact, we repeated the analysis in time-courses for various knockdowns, and found no cell-cycle signal in most cases (Figure 5F). A notable exception is the depletion of the essential chromatin remodeler Sth1, where we observed an opposite signature, i.e. the transcription of G1/S genes decrease later than other genes (Figures 5E, S5F), perhaps related to reports that interference with Sth1 causes a G2/M arrest (Angus-Hill et al., 2001; Cao et al., 1997; Du et al., 1998). Importantly, we did not observe gross changes to the cell-cycle as measured by DNA staining for at least two hours following auxin addition, except for a potential small decrease in the S-phase fraction (Figures S4E). However, after about

Since the G1 and S sets were the only ones to show this effect (in Xrn1 and Sth1), we were worried that this set of genes has some property that makes it more susceptible in this analysis. We therefore selected sets of genes with similar half-life and expression levels and repeated the analysis, but found no significant hits (Figures S5G-H), excluding the possibility of an analysis artifact emanating from extreme mRNA levels or degradation rates.

We conclude that the reduction in the transcription adaptation response in arrested cells, and the unique and transient cell cycle signature upon Xrn1 depletion in an unsynchronized population point to a link between the cell-cycle and the transcription adaptation response.

### Transient mRNA accumulation is recapitulated but dampened when upstream factors along the 5’-3’ branch are depleted

The cell-cycle link provided clues as to *when* and *how* cells sense that mRNA is out of balance, however, the question of *what* is sensed by cells to trigger the transcription adaptation response remains unclear. Xrn1’s function is largely attributed to 5’-3’ degradation after deadenylation and decapping (Figure 6A). We reasoned that we could pin-point the molecular constituent being sensed by perturbing other factors in the mRNA degradation network and studying the changes to the transcription adaptation response. Therefore we AID-tagged multiple components of this intricate network - Not1 (Cdc39), Dis3 (Rrp44), Rrp6, Rat1, Pop2, Pab1, Dcp2, Pan3, Pap2, Nrd1, Sen1, Nab3, and Ccr4 (Figure 6A). We performed a depletion time-course experiment in these strains and subjected hundreds of samples to cDTA-seq (Figure 6B).

**Figure 6.**
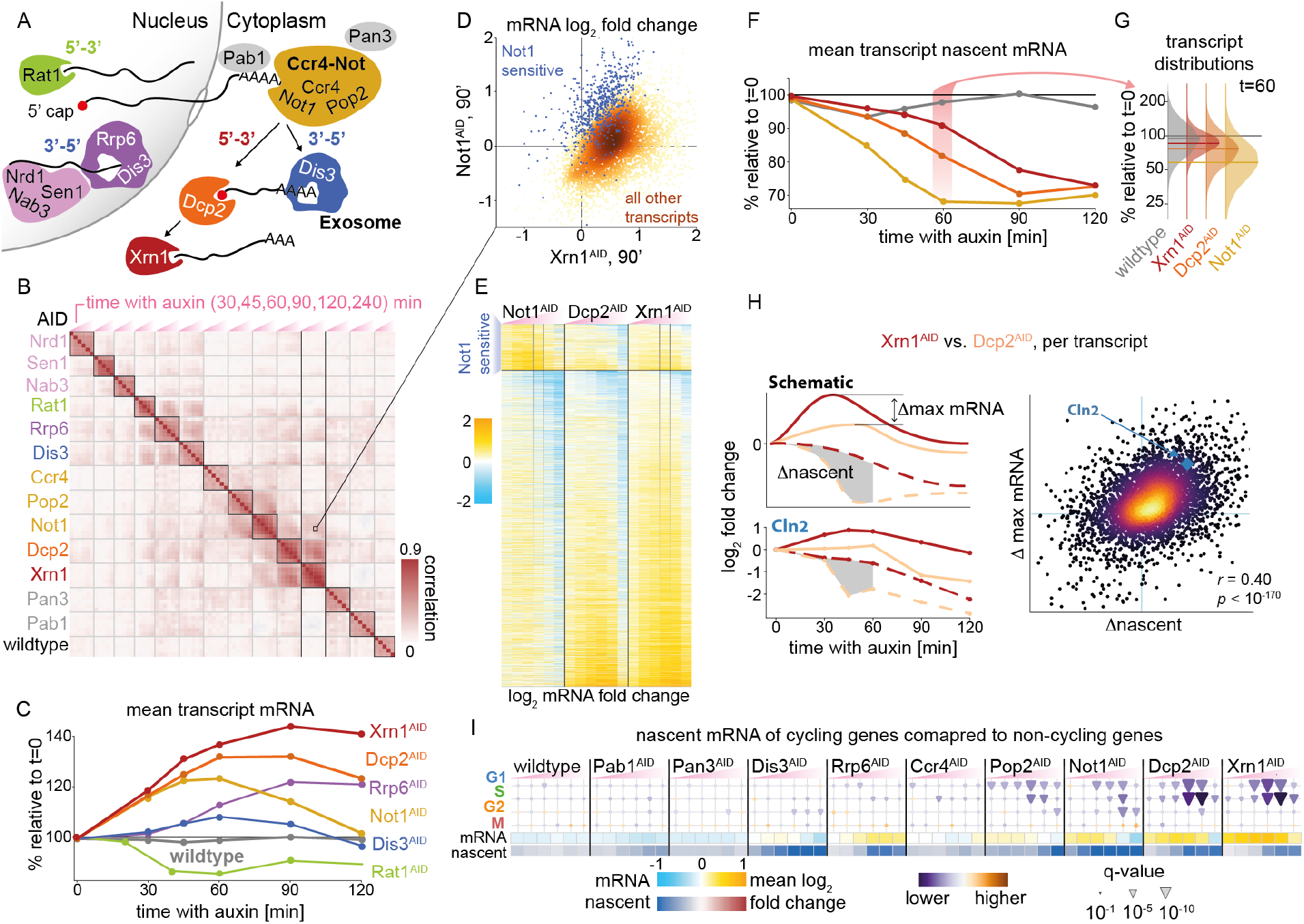
The transcription adaptation response is induced earlier when the 5’-3’ pathway is perturbed upstream. **A) AID-tagged proteins in their context.** Nuclear factors to the left - Rat1 is the nuclear 5’-3’ exoribonuclease, Nrd1, Sen1 and Nab3 survey aberrant transcripts and recruit the nuclear exosome (Dis3, Rrp6). Mature mRNA leaves the nucleus and will be deadenylated in the cytoplasm by the Ccr4-Not complex (Not1, Pop2, Ccr4). After deadenylation transcripts will continue to degrade 3’-5’ by the cytosolic exosome (Dis3), or 5’-3’ by Xrn1 after decapping by the DCP complex (Dcp2). Pab1 is the polyA binding protein, and Pan3 is part of an alternative deadenylation complex (Parker, 2012). **B) Depletion time course correlations.** Factors noted in (A) were subjected to a 4-hour depletion time course (Figure 3B), and cDTA-seq. The matrix summarizes the Pearson correlation between the log fold changes to transcripts relative to t=0 in each strain. Highlighted box corresponds to the relationship shown in (D). Xrn1 correlations are demarcated with black horizontal lines, significant correlations (Figure S6A) are elaborated in (C). **C) General mRNA response in time for selected factors.** Average changes to transcripts’ mRNA relative to t=0 (y-axis) along the time course (x-axis) for factors exhibiting significant correlation to Xrn1 (B, Figure S6A). **D) Correlation between Xrn1 and Not1 depletion.** Fold change relative to t=0 after 90 minutes in Xrn1^AID^ (x-axis) and Not1^AID^ (y-axis). Changes are generally correlated. Set of Not1-sensitive transcripts are marked in blue, other points are colored by density. Data corresponds to marked columns in (E) and marked box in (B). **E) mRNA changes upon interference to the 5’-3’ cytosolic pathway are correlated.** Changes to transcripts (rows) along the time-course (x-axis, 15, 30, 45, 60, 90, 120, 240 minutes) in three time-courses: upon depletion of Xrn1, Dcp2, and Not1. The mRNA log fold change relative to t=0 is color-coded. Transcripts are split into two sets - the upper set are transcripts that respond strongly to Not1 depletion (N = 672), and the lower are the rest of the transcripts (N = 4539). Rows in each set are sorted by the extreme point in a smoothed trajectory of the Xrn1 response. Highlighted columns correspond to x- and y-axis in (D). **F) The transcription adaptation response occurs earlier when the 5’-3’ pathway is perturbed upstream.** Average changes to transcripts’ nascent mRNA relative to t=0 (y-axis) along the depletion time course (x-axis). The highlighted 60 minutes time point is detailed in (F). **G) Transcript Variability in the response.** The distribution of changes in nascent mRNA per transcript relative to t=0 (y-axis) after 60 minutes of auxin in the three factors depleted in (F). The median of each distribution is denoted by a horizontal colored line. **H) Nascent mRNA differences explain mRNA differences between strains.** To compare the differences in response profiles between strains (in this case, comparing Xrn1 and Dcp2), we calculate the difference between the maximum observed change in mRNA per transcript (“Δmax”, y-axis in scatter) and between cumulative nascent trajectories (shaded grey area in examples, “Δnascent”, x-axis in scatter). We plot these statistics per transcript (dots in scatter, color denotes density) and found significant correlation (Pearson *r =* 0.4 *p* < 10^-170^). See Figure S6F for the same comparison between Dcp2 and Not1 (Pearson *r =* 0.31, *p* < 10^-98^). **I) Cell cycle signature only appears along the time course when depleting 5’-3’ factors.** We repeated the cell-cycle signature analysis (Figure 5G-H). Differences in log fold change to nascent mRNA between cycling gene sets (row) and non cycling genes denoted as colored triangles (purple, down - lower than cycling genes, orange, up - higher, size proportional to kolmogorov-smirnov q-value). Each triangle denotes the difference along a specific depletion time point (columns, x-axis time since auxin addition, same as in (B)). Bottom panels denote the average log fold change to mRNA and nascent mRNA in the same samples.

The changes in mRNA in this large dataset revealed the compartment, pathway, and protein-complex interactions between the depleted factors. For example, the effect of depletions of Rrp6, and Dis3 - subunits of the nuclear and core exosome respectively (Wasmuth et al., 2014) - are significantly correlated (as expected), but are also correlated with depletion of other nuclear proteins (Figure 6B). We wanted to further dissect the response to Xrn1 depletion, and focused on Xrn1-correlated factors - Rat1, Rrp6, Dis3, Not1, Dcp2 (Figure S6A). We examined the overall mRNA profile following the depletion of these five additional targets (Figure 6C), and it became clear that responses upon Dcp2 and Not1 depletion were significantly more similar to Xrn1 depletion than all other ones (Figure S6A).

Not1 and Dcp2 act upstream to Xrn1 in the 5’-3’ mRNA degradation pathway - Dcp2 is the catalytic component of the decapping complex, and Not1 (Cdc39) is the (essential) scaffold of the Ccr4-Not deadenylation complex. In all three depletion time-courses (Xrn1, Dcp2, Not1) we observed an accumulation of mRNA followed by a reduction in mRNA levels, consistent with a general feedback mechanism that is triggered when this pathway is perturbed (Figures 6C-E).

We compared the transcript profiles along the depletion time course (Figures 6D-E) and observed overall high correlation between the perturbations (10^-300^ < *p* < 10^-63^, Figure S6A). More specifically it seems that mRNA accumulation is most pronounced in the case of Xrn1 depletion, slightly dampened when Dcp2 is depleted, and further muted when Not1 is depleted.

Further examination of the data revealed a subset of ~13% of transcripts that were more sensitive to Not1 depletion (Figures 6D-E, S6B-C). A functional analysis of these transcripts shows a strong enrichment for transcripts of proteolysis-related genes (q < 10^-8^, Fig S6D). A link between Not1 and proteasome transcript regulation was reported in the literature, and recently, a co-translational complex assembly mechanism was suggested to be mediated by Not1 (Kandasamy et al., 2021; Laribee et al., 2007; Panasenko and Collart, 2011; Panasenko et al., 2019). The rapid and prominent increase in these transcripts, largely without a concomitant increase in transcription (Figure S6E) suggests that these transcripts are especially susceptible to Ccr4-Not-dependent degradation directly or via the 3’-5’ degradation pathway. These results expand the previously reported link between Ccr4-Not and post-transcriptional regulation of the proteasome. Importantly, even in the Ccr4-Not-sensitive transcript cluster, mRNA accumulation following depletion of Xrn1 and Dcp2 is consistent with the global response pattern (Figures 6D-E), suggesting that when Not1 is depleted a part of the observable increase in this cluster is due to accumulation emanating from the interference to the 5’-3’ degradation branch (Figures 6D, S6B-C).

These results demonstrate that mRNA generally accumulates in the same pattern when factors along the 5’-3’ degradation branch are perturbed, but the degree of accumulation depends on the specific element depleted (Not1 < Dcp2 < Xrn1).

### Upstream perturbations in the 5’-3’ degradation pathway result in earlier onset of the transcription adaptation response

To understand the apparent association between global mRNA accumulation profile and the 5’-3’ degradation pathway order, we excluded the Ccr4-Not sensitive genes from the analysis, and turned to examine nascent transcription along the depletion time-courses of these factors.

In all three cases we observed a reduction in nascent transcription, but strikingly, the timing order of the observed reduction recapitulated the observed order in the case of total mRNA, namely - Not1 caused the most immediate decline in nascent transcription, followed by Dcp2, and then by Xrn1 (*t_½_* of 33’, 52’, and 70’ respectively; Figure 6F). This result suggested that the difference in accumulated mRNA between the strains is due to earlier onset of the adaptation response in Not1 relative to Dcp2 and in Dcp2 relative to Xrn1.

While the reduction in nascent transcription is relatively synchronized across the genome, there is still significant transcript variation (Figure 6G). If the earlier reduction in nascent transcription explains the reduced degree of mRNA accumulation in Not1 relative to Dcp2 and relative to Xrn1, this relationship should also hold per transcript, namely, genes whose transcription is decreased faster in Dcp2 compared to Xrn1 should accumulate less mRNA in Dcp2 compared to Xrn1. To examine this hypothesis we calculated the difference between cumulative changes in nascent transcription during the accumulation phase (<60 minutes, “ΔNascent”), and the difference in maximal total mRNA during the accumulation and adaptation phase (<90 minutes, “max mRNA”, Figure 6H). We then compared these measures between the different strain pairs, and found a strong correlation (Fig 6H, *r* = 0.4, *p* < 10^-170^), i.e. genes with lower nascent transcription in Dcp2 compared to Xrn1 accumulated more mRNA in Xrn1 relative to Dcp2, as expected. This was also the case (albeit to a lesser degree) when we examined the differences between Dcp2 and Not1 (*r* = 0.31, *p* < 10^-98^, Figure S6F).

Finally, we reasoned that if the G1/S cell-cycle signature is an intrinsic part of the transcription adaptation response we observed in the case of Xrn1 depletion, and Not1 and Dcp2 are subject to essentially the same response, then the cell cycle signature for G1/S should be repeated in the case of Not1 and Dcp2. Indeed, when we performed the same enrichment analysis on these depletion time-courses (Fig 6I) we observed a strong and transient bias for G1/S genes to an earlier transcriptional shutdown only along the 5’-3’ branch (Xrn1, Dcp2, Not1, Pop2).

Taken together, these results point to a global mRNA accumulation pattern in response to a perturbation along the 5’-3’ degradation pathway. The degree of mRNA accumulation can be explained by the timing of the transcription adaptation response, and furthermore, the onset time of the response is earlier when upstream factors in the 5’-3’ pathway are perturbed.

## Discussion

We set out to study the mechanism of mRNA homeostasis that had been observed in various conditions and organisms. The literature surrounding the question of feedback mechanisms between mRNA degradation and transcription is largely based on steady-state measurements. As a baseline, when we examined Xrn1 knockout we recapitulated a previous observation of a genome-wide reduction in degradation and transcription rates resulting in unchanged global mRNA (Figure 2). We reasoned that to study a feedback mechanism, steady-state measurements can be insufficient and potentially lead to incorrect interpretations. Therefore we developed and applied a high-throughput sequencing-based implementation of the widely used cDTA technique (Figure 1). This allowed us to monitor total and nascent mRNA in detailed dynamic settings for hundreds of samples, resulting in the biggest resource of nascent transcription data in yeast to date. Applying cDTA-seq to cells undergoing rapid Xrn1 depletion (Figure 3A), we observed pronounced mRNA accumulation in virtually all transcripts, followed by a return to wildtype levels. The nascent mRNA profile provided by cDTA-seq revealed a striking reduction in nascent transcription roughly 60 minutes following Xrn1 depletion (Figure 4). An analysis of the time-course data also revealed a transient signature of G1 and S transcripts pointing to the cell-cycle as a potential component of the adaptation response (Figure 5). Finally, when we applied cDTA-seq to cells in which different RNA processing factors were depleted, we found that the response to Xrn1 depletion is not unique; detailed dynamic measurements revealed that the depletion of decapping and deadenylation factors results in a similar initial increase in mRNA levels. However, while the initial response was similar, the reduction in nascent transcription occurred earlier when upstream factors along the 5’-3’ degradation pathway were perturbed (Figure 6).

We focused on Xrn1 for two main reasons. First, it was implicated in mRNA homeostasis in two important but incongruent works by the Choder (Haimovich et al., 2013b) and Cramer (Sun et al., 2013) groups. Secondly, Xrn1 degrades a large proportion of mRNA molecules in eukaryotic cells (Geisler and Coller, 2012), so we expected a considerable response. Indeed, Xrn1 knockout exerts the most extreme alterations to mRNA profiles in published systemic knockout studies (Figure S2A). Despite our technique being more akin to the cDTA protocol used by the Cramer group (Figure 1G), our data supports the results from the Choder group, namely that mRNA levels are unchanged when cells lack Xrn1 and that degradation and transcription rates are significantly reduced (Figure 2).

To understand this homeostatic response we applied cDTA-seq to cells undergoing rapid Xrn1 depletion (Figure 3A). We observed pronounced and transient mRNA accumulation immediately after the perturbation. To our knowledge this is the first time that such a detailed view of a transient increase in mRNA levels is observed. Strikingly, in similar depletions of other factors (but not all, Figure 6C) cells revert to almost exactly the same mRNA levels they began with, suggesting some form of perfect adaptation taking place (Barkai and Leibler, 1997; Muzzey et al., 2009). Further dynamic data in different settings will be useful to model this possibility. Importantly, this response does not seem to be a normalization artifact as (i) it was measured in multiple different experimental batches and conditions (Figures 3, 5, and 6), and (ii) it was observed by single molecule FISH in four different transcripts (Figure 3). The observed reversion to normal levels was harder to verify by smFISH, as cells lacking Xrn1 accumulate deadneylated mRNA molecules (Hsu and Stevens, 1993; Wiener et al., 2021). This discrepancy makes the direct counting of non polyA molecules in single cells more challenging. Therefore, we used polyA probes and FACS measurements and observed a corresponding signal to the cDTA-seq signal (Figure S3D).

Having excluded normalization, increased growth, or induced degradation as possible explanations to the reduction in mRNA levels, we could explain the return to wildtype levels by a global reduction in nascent transcription roughly 60 minutes following Xrn1 depletion together with dilution by continued growth (Figure 4). The synchronous nature of the reduction strongly supports a general mechanism for transcriptional regulation, rather than the existence of multiple gene-specific feedback loops. This result is further supported by a recent report that uses aneuploid cells to distinguish between these two possibilities (García-Martínez et al., 2021b). Supplementing the observed reduction in nascent transcription, we also observed increased accumulation of cells in the G1 phase of the cell-cycle, but only after 2.5 hours (Figure 5E). While the observed delay in G1 cannot explain the reduction in nascent transcription after 60 minutes, it probably is a factor in the observed reversion to basal mRNA levels, as G1 cells have roughly half the amount of mRNA compared to G2 cells (Voichek et al., 2016), which would, on average, reduce the amount of mRNA per cell in the population.

Importantly, the observed delay in nascent transcription reduction argues against direct involvement of Xrn1 in this transcriptional reprogramming, as Xrn1 is degraded rapidly from cells (Figure 3A), but the transcriptional response occurs an hour later (Figure 4). Conversely, when we depleted Rat1 (the 5’-3’ exonuclease operating in the nucleus that is involved in transcription termination (Kim et al., 2004)) we observed an immediate reduction in transcription (e.g. compare nascent transcription in Figure 5E). Alternatively, Xrn1 was hypothesized to indirectly affect global transcription through post-transcriptional regulation of Nrg1(Sun et al., 2013). We therefore repeated the Xrn1 depletion time course experiment in ΔNrg1 and ΔNrg2 strains but we did not observe any difference in the total or nascent mRNA dynamics (not shown), arguing against Nrg1/2 involvement in the feedback.

Having ruled out the two main proposed mechanisms by which Xrn1 affects transcription we looked for alternative explanations. The time scale of the delay (60-90 minutes) led us to examine the cell cycle as a potential component of the homeostatic mechanism. When we prevented cells from iterating through the cell cycle only a modest decrease in transcription was observed following Xrn1 depletion (Figures 5B). Extending this result, when we tested the response in asynchronous populations we found a clear cell cycle signature only in the case of Xrn1 depletion and other perturbations to the 5’-3’ mRNA degradation pathway (Figures 5F, 6I). Even though both lines of evidence are circumstantial, they point to a role for the cell cycle in the observed feedback. The observed G and S signature in the unsynchronized population and the delayed reduction in nascent transcription, suggest a sensing mechanism that is triggered in a specific stage of the cell cycle, after a period of time has passed since Xrn1 depletion (Figure 7A).

**Figure 7.**
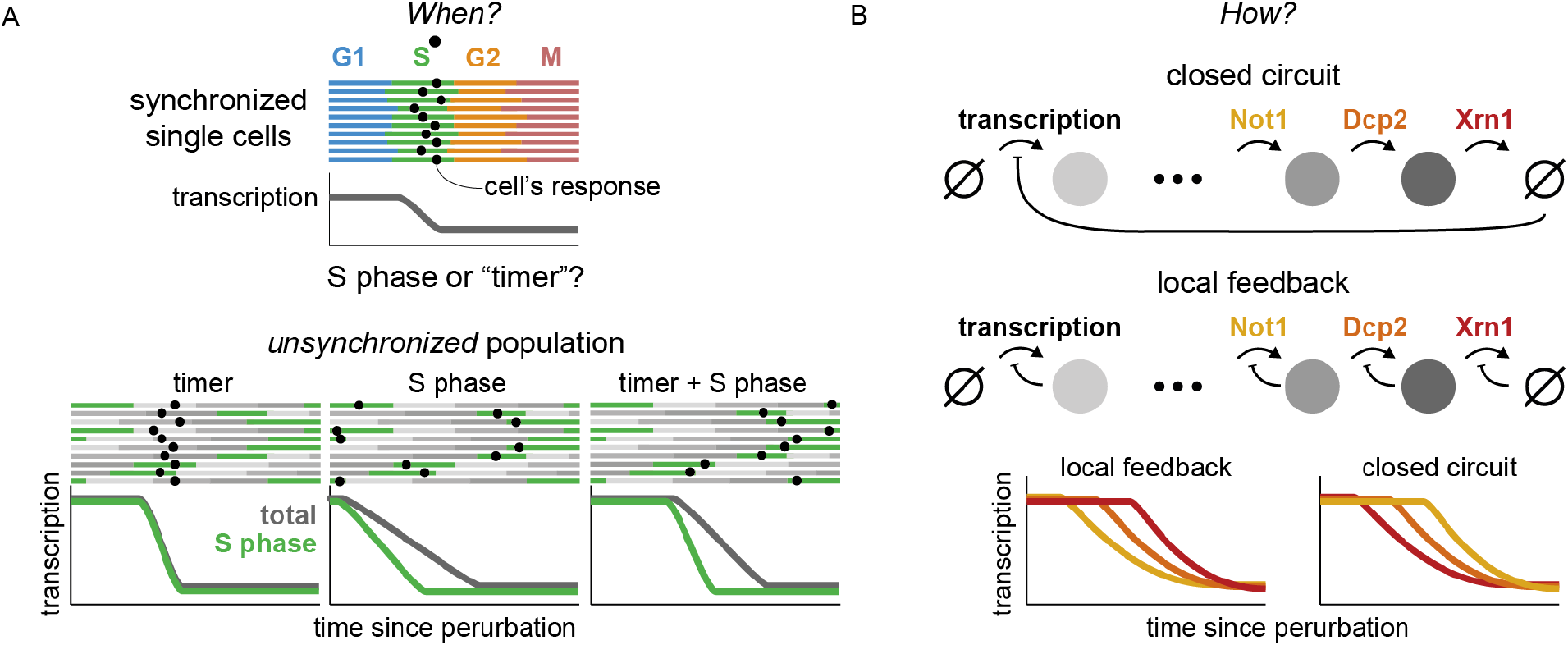
Dynamic measurements distinguish between different feedback models. **A) Cell cycle data suggests a cell-cycle coupled timer sensing mechanism.** In a synchronized population (top) a delayed transcriptional response can be explained by a “timer” measuring a cellular element accumulating since Xrn1 depletion, or by a specific cell cycle phase (S-phase as an example) that monitors e.g. Xrn1 levels. However, dynamic measurements from an unsynchronized population can be used to distinguish these models by the differences between genes expressed along the cell cycle (in this case in S-phase, green) to non-cycling genes (dark grey). Our data suggest a model involving the cell cycle (G1/S) and a timer mechanism. **B) Temporal offsets between different perturbations suggest local rather than a closed circuit feedback mechanism.** Two toy models that result in global mRNA feedback are presented. In the closed circuit model transcription is coupled to degradation directly, while the local feedback model assumes that each stage self-regulates. While the steady-state behavior of both models will be similar, dynamic measurements can be used to distinguish the two by the delay in the propagation of the interference back to transcription. Our data are consistent with a local feedback model.

Cell cycle checkpoints are known to monitor cell size, nutrients, DNA damage, and proper chromosomal and cellular geometry (Barnum and O’Connell, 2014), and it is possible that the accumulation of mRNA (or some other cellular element) feeds back into one of these sensors or to a yet undescribed checkpoint. Furthermore, a cell-cycle coupled mechanism for regulating global transcription during the S phase was previously described (Padovan-Merhar et al., 2015; Voichek et al., 2016), making the cell cycle an attractive candidate for implementing the hypothesized sensing-acting mechanism. Since much of the molecular details of these checkpoints are known, testing the link between the transcriptional adaptation we observed in response to mRNA accumulation and the cell cycle should be feasible.

To further dissect the response, we aimed to pinpoint the molecular species being sensed by cells. We expanded our experiments to include the depletion of dozens of RNA-related factors (shown in Figure 6A, and others, not shown). This expansive view revealed various response dynamics (Figures 6B-C) and will hopefully prove useful to understand different phenomena related to mRNA processing. Focusing on the adaptation response we observed in the wake of Xrn1 depletion, it was clear that Dcp2 and Not1 depletion elicited similar responses (Figures 6C, S6A). Strikingly, the mRNA response in all three cases began essentially the same, but seemed to be attenuated in the order of the factor along the 5’-3’ mRNA degradation pathway (Not1 < Dcp2 < Xrn1). Consistently, we observed a delayed reduction in nascent transcription, that also followed the order of the perturbation along the 5’-3’ degradation pathway (Figures 6A, 6F), parsimoniously explaining the reduced accumulation of mRNA (Not1 < Dcp2 < Xrn1). We interpret these results to suggest that cells do not monitor accumulated deadenylated and decapped mRNA, or downstream byproducts such as p-bodies(Sheth and Parker, 2003), for if this was the case, transcription inhibition would ensue earlier in the Xrn1 depletion time course (Figure 7B). Extending the logic of this argument, the agreement between the 5’-3’ mRNA degradation pathway order and the onset time of the transcription adaptation response suggests that cells monitor a precursor that accumulates upstream of this pathway.

This model, when extrapolated, suggests local feedback along the lifecycle of the mRNA, rather than a closed circuit feedback mechanism (Figure 7B). Specifically, the model posits that accumulation of decapped mRNA inhibits decapping by Dcp2, and that subsequent accumulated capped mRNA inhibits deadenylation by Ccr4-Not, which will explain the timing offsets we observed. There are several interesting options upon consideration of this prediction: (i) It is possible that 5-AMP released by Ccr4-Not deadenylation (Tucker et al., 2001) is important for proper cellular metabolism in general or for DNA replication specifically. A rough calculation revealed that polyA-bound adenosine is within one order of magnitude of the amount of free ATP in cells (Koç et al., 2004). (ii) As previously suggested, polyA binding protein (Pab1) could be involved in the sensing mechanism (Gilbertson et al., 2018; Kumar et al., 2011). Notably, the Pab1 depletion data we presented here (Figure 6) argue against this hypothesis, but it requires further scrutiny, and we have not directly tested the nuclear analog - Nab2(Schmid et al., 2015). (iii) In a similar vein, an imbalance in mRNA nuclear export might be caused by polyA mRNA accumulation, which in turn causes an accumulation of nuclear mRNA that was recently suggested to directly affect transcriptional throughput (Berry et al., 2021; Henninger et al., 2021). (iv) Ribosomes are essential for growth and proliferation, and free ribosomes were suggested to play an important role in this regulation (Metzl-Raz et al., 2017; Weiße et al., 2015). It is possible that cells monitor the amount of free ribosomes which is presumably reduced as poly-A mRNA accumulates. (v) Alternatively, an imbalance between ribosomal proteins and rRNA can cause downstream nuclear dysfunction and cell-cycle progression defects (Gómez-Herreros et al., 2013). Such an imbalance can arise from over-production of ribosomal proteins due to mRNA accumulation. (vi) Last, the Ccr4-Not complex has been suggested to be a hub affecting transcription, translation, and degradation (Collart, 2016). It is possible that a functional diversion of the Ccr4-Not complex itself due to accumulated mRNA causes the transcriptional response.

Taken together, our results suggest a model (Figure 7) in which some input to the 5’-3’ mRNA degradation pathway in cells is being monitored and once a critical threshold is met at a certain phase of the cell cycle, an adaptive transcriptional response ensues, allowing cells to reestablish proper mRNA levels. More broadly, while mRNA homeostasis observations and functional experiments were mostly conducted in yeast, there is a growing body of literature that suggests this phenomenon is of a more general nature (Berry et al., 2021; Helenius et al., 2011; Kumar et al., 2011; Slobodin et al., 2020). Notably, the factors we identified here along the 5’-3’ mRNA degradation pathway are highly conserved. A better understanding of the feedback mechanism employed by cells will likely have implications beyond yeast and will be relevant for critical processes in health and disease such as proliferation, apoptosis, and viral immune response (Duncan-Lewis et al., 2021). The data we presented here argue that detailed dynamic measurements are crucial in the search for a mechanistic understanding of this process.

## Acknowledgements

We thank Michal Chappleboim and Miri Carmi for their help with single-molecule FISH. We thank Ronen Sadeh, Jenia Gutin, Michal Rabani, Felix Jonas, and Naama Barkai for their comments on the manuscript. This work was supported in part by the Israel Science Foundation. AC was an Azrieli scholar and would like to thank the Azrieli Foundation for their support.

## Methods

### cDTA-Seq

#### Sample preparation

To compare mRNA levels between samples in each experiment, *Saccharomyces cerevisiae* (SC) cells were quantified, and fixed in frozen methanol that was pre-spiked with *Kluyveromyces lactis* (KL) yeast cells. For each experimental batch, KL cells were grown to log-phase, fixed in frozen methanol and the KL-spiked methanol was equally pre-distributed in deep well plates in relatively large volumes to avoid pipetting issues. The plate with pre-spiked methanol was kept at −80C until the fixation of SC samples. Immediately prior to SC fixation, cell density was measured by optical density at 600 nm and the rest of the sample was metabolically labeled by 4tU for several minutes. This was performed simultaneously to the entire plate, after which cells were immediately transferred to fixation in pre-frozen and pre-KL-spiked methanol (600 μl pre-spiked frozen methanol to 500 μl SC cells). Samples were then kept in −80°C for varying duration ranging from days to several weeks.

#### RNA purification

Cells fixed in frozen methanol were washed twice in ddw and RNA purification was performed as previously described (Dye et al., 2005) with minor modifications. Briefly, RNA was released from the cells by digestion with Proteinase K (Epicenter) in the presence of 1% SDS at 70°C. Cell debris and proteins were precipitated by centrifugation in the presence of potassium acetate. RNA was then purified from the supernatant using nucleic acid binding plates (96-well, 800 μl UNIFILTER Microplate, GE Healthcare) in the presence of 0.1 mM DTT, eluted in 1 mM DTT and stored at −80°C.

#### Metabolic labeling, and adaptation of SLAM-seq to yeast

Metabolic labeling of new RNA molecules was done as previously described (Herzog et al., 2017; Voichek et al., 2016). Briefly, 4-thiouracil (4tU, Sigma) was dissolved in NaOH and added to cells at final concentration of 5 mM 4tU for the indicated times (6-10 minutes). To avoid pH change as a result of NaOH addition, MES buffer was added to the media prior to growth. RNA purification was performed as described above. Total RNA was subjected to thiol(SH)-linked alkylation by iodoacetamide (Sigma, 10 mM) at 50^0^C for 15 minutes, the reaction was stopped with 20 mM DTT. RNA was purified using nucleic acid binding plates (96-well, 800 μl UNIFILTER Microplate, GE Healthcare) and was stored with RNAse-inhibitor at −80°C.

#### polyA RNA-seq library preparation

Library preparation was done as previously described (Klein-Brill et al., 2019). Total RNA was incubated with oligo-dT reverse transcription primers with a 7 bp barcode and 8 bp UMI (Unique Molecular Identifier) at 72°C for 3 minutes and transferred immediately to ice. RT reaction was performed with SmartScribe enzyme (SMARTScribe Reverse Transcriptase, Clontech) at 42°C for one hour followed by enzyme inactivation at 70°C for 15 minutes. Barcoded cDNA samples were then pooled and purified using SPRI beads x1.2 (Agencourt AMPure XP, Beckman Coulter). DNA-RNA hybrids were tagmented using Tn5 transposase (loaded with oligos Tn5MEDS-A, Table S2) and 0.2% SDS was added to strip off the Tn5 from the DNA (Picelli et al., 2014), followed by a SPRI x2 cleanup. Barcoded Illumina adaptor sequences (Table S2) were added to the tagmented DNA by PCR (KAPA HiFi HotStart ReadyMix, Kapa Biosystems, 12 cycles). And the DNA was cleaned with an x0.8 SPRI procedure. Libraries were sequenced using Illumina NextSeq-500 sequencer.

#### Data processing

##### Demultiplexing

Pooled libraries were demultiplexed using Illumina’s bcl2fastq (version 2.20.0). Internal (sample) barcodes were demultiplexed with an awk command, not allowing any barcode errors.

##### Alignment

Prior to read alignment we prepared several versions of the SC and KL genomes. First, we converted both genomes to accommodate alignments of partially (T→C)-converted reads. This is achieved by converting all the observed Ts in the genome to Cs per strand. This results in an ACG-only genome with one contig per reference strand. In addition, We also generated a redacted version of the KL genome in which any 18-mer that is found in the SC genome was removed. This procedure removed only ~3.4% of the KL genome while increasing the proportion of reads that uniquely align to the KL genome from ~2.5% to ~99% (i.e. most reads arise from regions that are shared between the genomes).

polyA stretches were removed from read 3’ ends, and reads with more than 25 bases remaining were aligned in several different ways (using default bowtie2 settings for single end alignment):

1. SC genome without ACG conversion
2. SC genome with ACG conversion
3. KL genome without ACG conversion
4. KL genome with ACG conversion
5. Redacted KL genome without ACG conversion

These allow for quality control measures to be calculated per sample. However, for downstream analysis, only the (2) and (5) alignments are used. Alignment after ACG conversion was performed by converting observed Ts to Cs in reads and aligning them against the converted genome. Following alignment to the converted genome by bowtie2, a dedicated script converts reads back to the original reference coordinates and strand, and marks any sequence discrepancies between the original observed sequence and the reference sequence per read.

##### Read filtration and UMI handling

Reads in all sequencing runs had lengths between 44-46, and we discarded reads with more than 30 observed Ts (binomial probability < 4e-7, observed fraction ~ 4e-3). For typical analyses of mRNA, we only considered reads that were aligned less than five times with at most 3 errors excluding T→C conversions. Out of these alignments, only the best one was reported. Reads were de-duplicated by grouping according to (chromosome, strand, position, and UMI), and in each such group, the read with least amount of deviations from the reference was selected. Libraries had an average of 1.2-1.5 reads per UMI.

##### Read statistics

Filtered and deduplicated alignments were assigned to transcripts by their intersection (Stovner and Sætrom, 2020) with a window of 300 bp upstream and 100 bp downstream of previously reported transcription termination site annotations (Weiner et al., 2015). Read statistics were then collected on individual transcripts, groups of transcripts, or the whole transcriptome in the form of a table with the number of reads for each combination of observed Ts and observed T→C vonersions.

##### Relative mRNA level estimation

To estimate the relative amount of mRNA in each sample we consider only the reads that were aligned to annotated TTS regions (see above). We divide this number by the sample OD, and by the total number of reads that were aligned to the KL redacted genome. While this procedure removes most of the variance between replicates, in time course experiments we also smooth these estimates with a savitzky-golay filter (3rd degree) spanning a 120 minute window.

##### Binomial Mixture Model

Similar to (Jürges et al., 2018), we assume that reads arrive from a mixture of two sets - old transcripts that were transcribed prior to the labeling period, and new transcripts that were generated after the labeling period began. To estimate the typical molecule half-life, we are interested in the relative size of each of these sets. This proportion can be denoted with a single parameter - the nascent fraction - *p_n_*. The model stipulates that if a molecule arrives from the “old” set then we expect T→C conversions at a certain rate - *ε*, if however, the molecule is from the nascent set, then we should observe conversions at a higher rate - *ξ*. In either case, if a read has X observed Ts and Y observed conversions, assuming a uniform rate along the read, the probability of observing Y conversions should be binomially distributed:

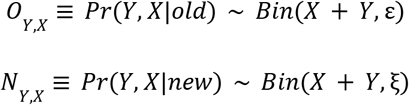

Therefore the overall probability of observing Y, X is:

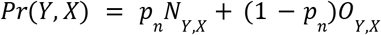

And more generally, the likelihood of a collection of reads, R:

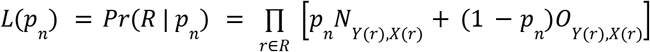

Thus, assuming the error and conversion rates are global (Figure S1E), the likelihood is a function of a single parameter - *p_n_* - ranging from 0 to 1, and its maximum can be efficiently calculated given the other model parameters (supplementary note).

##### Steady-state model

We assume the following dynamic system:

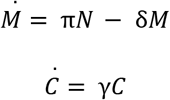

Where *M* is the total amount of mRNA, C is the number of cells, π is gene-specific transcription rate per cell, *δ* is the gene-specific degradation rate, and *γ* is the growth rate. Redefining 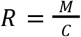, i.e. mRNA/cell, the system simplifies to:

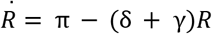

We also assume that 4tU labeling does not disturb cells from their exponential growth pseudo-steady-state (Figures S1G-H), in which case the steady state value of the system is given by:

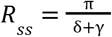

Additionally, the differential equation yields the following dynamics for nascent transcripts (*N*):

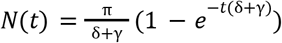

Thus, the proportion of nascent transcripts, *p_n_*, should evolve with labeling time, *t*, as follows:

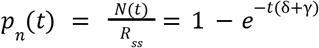

##### Half-life estimation

Briefly, we fit/measure several global parameters (detailed description is given in the supplementary material): incorporation probability (*ξ*) error probability (*ε*) conversion lag time (*t*_0_), and growth rate (*γ*). Once these are determined, the only gene-specific parameter is the degradation rate (*δ*), which can be calculated by fitting the BMM model to the data to obtain *p_n_*, and inverting the last equation:

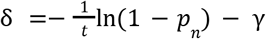

Which can be further transformed to a half-life (ln(2)/*δ*).

Detailed specific analyses to generate all the figures and data are provided as python notebooks and further explained in the supplementary material.

### Auxin-induced degradation

In all experiments, yeast cells were grown in YPD supplemented with 10 mM MES buffer at 30 degrees overnight. 60 minutes prior to the beginning of the time course, samples with OD ranging from 0.3 to 0.6 were split to deep well plates and grown at 25 degrees with constant pipette mixing during the time course. At indicated times auxin (3-indolo acetic acid, Sigma) was added at a final concentration of 1-2.5 mM (table S4). Auxin stock was dissolved in DMSO to 2.5M, and diluted 1:1 in 1M NaOH before being added to samples to prevent sedimentation in the aqueous medium. When mock treatment was appropriate, the same amount of DMSO and NaOH was added to control samples.

### cDTA-seq calibrations

#### Spike-in titration

To verify we can robustly quantify and compare mRNA levels per cell between samples, we need two points of reference - the first is the amount of sample mRNA compared to an absolute standard, and the second is the amount of cells from which the mRNA was extracted. As a standard for mRNA levels we spike-in fixed amounts of exogenous cells to each sample. To evaluate this strategy, we performed a titration of spike-in cells (*Kluyveromyces lactis* - KL) in the 1%-5% range into a *Saccharomyces cerevisiae* (SC) sample. SC and KL cells were grown to log-phase (od 0.5) and fixed separately in cold methanol (7.5 ml cells in 9 ml methanol). Fixed cells were mixed at varying ratios (1-5% KL) and were kept at −80°C. RNA purification and polyA RNA-seq libraries were prepared as described above.

#### Transcription inhibition

Thiolutin (Sigma) was dissolved in dimethyl sulfoxide and added to cells at final concentration of 3 μg/ml for 15 minutes.

#### DNA-seq validation

Cell quantification is typically done manually(Sun et al., 2012) but this procedure is prohibitive when analyzing dozens and hundreds of samples. In many yeast experimental settings the optical density (OD) of the culture is a good proxy to the cell counts per unit volume. However, there are concerns that different genetic backgrounds on cell states can affect the OD/cell ratio(Kokina et al., 2014; Stevenson et al., 2016). We verified that our OD measures correspond to cell counts by comparing the DNA content extracted from 93 samples from various genetic backgrounds to their OD. To estimate the ratio between the number of SC and KL cells we extracted nucleic acids from various samples that underwent cDTA-seq and were therefore spiked-in with KL cells. These samples included various AID-tagged proteins (Dcp2, Xrn1, Rat1, Fcp1, Sth1, Med14, Pop2, Spt6) grown overnight with varying degrees of auxin, resulting in a wide range of growth rates (1.5-4 hour doubling time) and morphological phenotypes (e.g. Xrn1-depleted cells are larger). When we compared the ratio between SC DNA and KL DNA in each sample to the OD of that sample we observed high correlation (Figure S1B).

To prepare DNA, RNase (Sigma, 11119915001) was added to nucleic acid extracted as mentioned in the cDTA-seq protocol (0.1 μl RNase to 100 ng RNA in a reaction volume of 20 μl) and incubated at 37°C for 30 minutes. Remaining material was tagmented using Tn5 transposase (loaded with oligos Tn5MEDS-A and Tn5MEDS-B, Table S2) and 0.2% SDS was added to strip off the Tn5 from the DNA (Picelli et al., 2014). Barcoded Illumina adaptor sequences (Table S2) were added to the tagmented DNA by PCR (KAPA HiFi HotStart ReadyMix, Kapa Biosystems, 20 cycles). And the DNA was cleaned with a x0.9 SPRI procedure. Libraries were sequenced using Illumina NextSeq-500 sequencer.

Reads aligned to the SC genome were manually inspected to have uniform genomic distribution (with exceptions in the rDNA locus, transposable elements, etc.). Reads were also aligned to a redacted SC genome (removing any 20-kmer found in KL) and to a similarly redacted KL genome to obtain estimates to the amount of DNA from each organism in the sample. The ratio between these numbers was compared to the OD of each sample (both measures were normalized to their median for visualization in Figure S1B). Outliers in the scatter (Figure 1C) were not of a specific strain, batch, or condition, pointing to technical measurement issues (probably in the OD), rather than inherent biases in the technique.

### Western blots

Yeast lysates were prepared as previously described(von der Haar, 2007) and protein were analysed using standard western blotting procedures with anti-FLAG M2 (sigma F1804), and anti-Myc (sigma M4439 clone 9E10).

### Growth assays and OD measurements

Optical density was collected using a Tecan Infinite F200 for 96 sample plates. Each sample was measured at five different points and the median value was used as the OD measure. A background level was subtracted from measured OD.

Growth assays were conducted using a Tecan Freedom Evo 2000 liquid handling station. The 200 μl 96 sample plate was incubated at 30°C, for >24 h with an automatic scheduled OD measurement in the Tecan Infinite F200 executed every hour ((Gutin et al., 2015).

A linear fit was applied to each consecutive set of 10 data points, and the minimal doubling time was determined by the fit with the highest slope among the fits with a significant (alpha 0.05) bonferroni-corrected p-value.

### Microscopy

Samples along the depletion time course were fixed in 1% formaldehyde for 15 min, and quenched in 125mM Gly. Bright field images were automatically collected in multiple fields per sample along an 8 μm z-stack (1μm step size) using a Scan^R system (Olympus). Cells were segmented using a freely available yeast segmentation software (Lu et al., 2019).

### Single molecule FISH

#### Sample preparation

In the first experiment, 120 ml Spt6^AID^ and 120 ml Xrn1^AID^ cells were grown to midlog, and split. Half of the culture was supplemented with auxin (final conc. 2.5 mM) and half with mock treatment (50% DMSO, 0.5M NaOH at equal volume). After one hour of incubation at room temperature samples were fixed in 5% formaldehyde and prepared as previously described(Rahman and Zenklusen, 2013) with fluorophore-conjugated (TAMRA or CAL Fluor Red 590) tiling probes for Msn2, Cln2, Cln3, and Suc2 (Biosearch technologies) were a gift from Naama Barkai and Jeffery Gerst (Msn2, Cln2, Cln3 sequences detailed in Table S3, Suc2 as in (Cohen-Zontag et al., 2019)).

In the second experiment, 40 ml of Xrn1^AID^ cells were grown to midlog and split to four samples. Samples were supplemented with auxin so to have a 240, 120, 60, and 0 time points in an auxin time course. Samples were fixed and prepared with the same probes for Msn2, Cln2, Suc2, and with a TAMRA-20dT probe ordered from IDT.

#### Microscopy

Images were acquired with a 100× 1.4 oil UPLSAPO objective, using an Olympus IX83 based Live-Imaging system equipped with CSU-W1 spinning disc (sCMOS digital Scientific Grade Camera 4.2 MPixel, Oxford Instruments, Abingdon, UK). For each sample, at least 4 different positions were chosen. In each position, three-channel Z-stacks images were taken with a step size of 200 nm for a total of >8 μm: bright-field image, 488 nm laser with 100 mW; DAPI image, 405 nm laser with 120 mW and exposure time of 250 ms; mRNA image, 561 nm laser with 100 mW and exposure time of 1,000 ms. Each z-plane image was of size 2,048 × 2,048 pixels.

#### Image analysis

Nuclei and single molecules were segmented from the DAPI channel using a MATLAB script that uses basic image processing steps (erosion/dilation/convolution) to account for uneven illumination. Segmented nuclei were filtered by size and manually inspected. Single molecule counts were obtained by using a custom-made MATLAB software (Raj et al., 2008).

#### polyA FACS analysis

At least 7,500 valid events were collected per sample on an Amnis CellStream high throughput flow cytometer after excitation with a 561 nm laser and acquisition with a 561-583 nm filter.

### DNA staining

Wildtype, Xrn1^AID^, and Sth1^AID^ cells were split and exposed to auxin along a time course of 4.5 hours. At the end of the time course, cells were collected into pre-frozen 100% EtOH (final 70%). Cells were washed in 50mM Tris-HCL pH 8 (Sigma), incubated with RNaseA (Sigma R4875, final 1mg/ml in 50 mM Tris-HCL pH8), Proteinase K (Sigma P2308, final 2.5 mg/ml in ddw), and with SYBR green ([Molecular Probes](http://www.probes.com/servlets/product?item=7563) S7567, diluted 1: 1000 in 10mM Tris-HCl pH8, 1mM EDTA pH8). Following an additional wash, cells were briefly sonicated and analyzed by FACS with a 525-centered filter (BD LSRII system, BD Biosciences). Cell-cycle phases were determined by fitting the DNA content distribution with a 3-component GMM.

### Strains and plasmids

*Saccharomyces cerevisiae* yeast strains used in this study are provided in Table S1.

Yeast strains were generated using the LiAc transformation method (Gietz et al., 1995). Auxin inducible degradation domain was PCR-amplified from plasmid pNat-AID*-9MYC or pHyg-AID*-6FLAG using matching primers (table S2) and introduced into TIR1 expressing cells immediately before the target gene stop codon (plasmids pHyg-AID*-6FLAG and pNat-AID*-9MYC and TIR1 cells are a gift from Ulrich lab (Morawska and Ulrich, 2013).

Fresh KOs (aa72, aa73) were prepared using plasmids pNat-AID*-9MYC or pHyg-AID*-6FLAG, and oligos(xrn1_ko_F, xrn1-deg-R) (table S2)

Oligos used in this study are listed in Table S2.

## References

Anderson, J.S., and Parker, R.P. (1998). The 3’ to 5’ degradation of yeast mRNAs is a general mechanism for mRNA turnover that requires the SKI2 DEVH box protein and 3’ to 5’ exonucleases of the exosome complex. EMBO J. 17, 1497–1506.

Angus-Hill, M.L., Schlichter, A., Roberts, D., Erdjument-Bromage, H., Tempst, P., and Cairns, B.R. (2001). A Rsc3/Rsc30 zinc cluster dimer reveals novel roles for the chromatin remodeler RSC in gene expression and cell cycle control. Mol. Cell 7, 741–751.

Baptista, T., Grünberg, S., Minoungou, N., Koster, M.J.E., Timmers, H.T.M., Hahn, S., Devys, D., and Tora, L. (2017). SAGA Is a General Cofactor for RNA Polymerase II Transcription. Mol. Cell 68, 130–143.e5.

Barkai, N., and Leibler, S. (1997). Robustness in simple biochemical networks. Nature 387, 913–917.

Barnum, K.J., and O’Connell, M.J. (2014). Cell Cycle Regulation by Checkpoints. Methods in Molecular Biology 29–40.

Baudrimont, A., Voegeli, S., Viloria, E.C., Stritt, F., Lenon, M., Wada, T., Jaquet, V., and Becskei, A. (2017). Multiplexed gene control reveals rapid mRNA turnover. Sci Adv 3, e1700006.

Begley, V., Jordán-Pla, A., Peñate, X., Garrido-Godino, A.I., Challal, D., Cuevas-Bermúdez, A., Mitjavila, A., Barucco, M., Gutiérrez, G., Singh, A., et al. (2021). Xrn1 influence on gene transcription results from the combination of general effects on elongating RNA pol II and gene-specific chromatin configuration. RNA Biol. 18, 1310–1323.

Berry, S., Müller, M., and Pelkmans, L. (2021). Nuclear RNA concentration coordinates RNA production with cell size in human cells. bioRxiv.

Blasco-Moreno, B., de Campos-Mata, L., Böttcher, R., García-Martínez, J., Jungfleisch, J., Nedialkova, D.D., Chattopadhyay, S., Gas, M.-E., Oliva, B., Pérez-Ortín, J.E., et al. (2019). The exonuclease Xrn1 activates transcription and translation of mRNAs encoding membrane proteins. Nat. Commun. 10, 1298.

Braun, K.A., and Young, E.T. (2014). Coupling mRNA Synthesis and Decay. Molecular and Cellular Biology 34, 4078–4087.

Bregman, A., Avraham-Kelbert, M., Barkai, O., Duek, L., Guterman, A., and Choder, M. (2011). Promoter elements regulate cytoplasmic mRNA decay. Cell 147, 1473–1483.

Cao, Y., Cairns, B.R., Kornberg, R.D., and Laurent, B.C. (1997). Sfh1p, a component of a novel chromatin-remodeling complex, is required for cell cycle progression. Mol. Cell. Biol. 17, 3323–3334.

Celik, A., Baker, R., He, F., and Jacobson, A. (2017). High-resolution profiling of NMD targets in yeast reveals translational fidelity as a basis for substrate selection. RNA 23, 735–748.

Chan, L.Y., Mugler, C.F., Heinrich, S., Vallotton, P., and Weis, K. (2018). Non-invasive measurement of mRNA decay reveals translation initiation as the major determinant of mRNA stability. Elife 7.

Chattopadhyay, S., Garcia-Martinez, J., Haimovich, G., Khwaja, A., Barkai, O., Ghosh, A., Chuarzman, S.G., Schuldiner, M., Elran, R., Rosenberg, M., et al. RNA-controlled nucleocytoplasmic shuttling of mRNA decay factors regulates mRNA synthesis and initiates a novel mRNA decay pathway.

Cohen-Zontag, O., Baez, C., Lim, L.Q.J., Olender, T., Schirman, D., Dahary, D., Pilpel, Y., and Gerst, J.E. (2019). A secretion-enhancing cis regulatory targeting element (SECReTE) involved in mRNA localization and protein synthesis. PLoS Genet. 15, e1008248.

Collart, M.A. (2016). The Ccr4-Not complex is a key regulator of eukaryotic gene expression. Wiley Interdisciplinary Reviews: RNA 7, 438–454.

Collart, M.A., and Struhl, K. (1994). NOT1(CDC39), NOT2(CDC36), NOT3, and NOT4 encode a global-negative regulator of transcription that differentially affects TATA-element utilization. Genes Dev. 8, 525–537.

Decker, C.J., and Parker, R. (1993). A turnover pathway for both stable and unstable mRNAs in yeast: evidence for a requirement for deadenylation. Genes Dev. 7, 1632–1643.

van Dijk, E.L., Chen, C.L., d’Aubenton-Carafa, Y., Gourvennec, S., Kwapisz, M., Roche, V., Bertrand, C., Silvain, M., Legoix-Né, P., Loeillet, S., et al. (2011). XUTs are a class of Xrn1-sensitive antisense regulatory non-coding RNA in yeast. Nature 475, 114–117.

Dori-Bachash, M., Shalem, O., Manor, Y.S., Pilpel, Y., and Tirosh, I. (2012). Widespread promoter-mediated coordination of transcription and mRNA degradation. Genome Biol. 13, R114.

Du, J., Nasir, I., Benton, B.K., Kladde, M.P., and Laurent, B.C. (1998). Sth1p, a Saccharomyces cerevisiae Snf2p/Swi2p Homolog, Is an Essential ATPase in RSC and Differs From Snf/Swi in Its Interactions With Histones and Chromatin-Associated Proteins. Genetics 150, 987–1005.

Duek, L., Barkai, O., Elran, R., Adawi, I., and Choder, M. (2018). Dissociation of Rpb4 from RNA polymerase II is important for yeast functionality. PLoS One 13, e0206161.

Duncan-Lewis, C., Hartenian, E., King, V., and Glaunsinger, B.A. (2021). Cytoplasmic mRNA decayre presses RNA polymerase II transcription during early apoptosis. Elife 10.

Dunckley, T., and Parker, R. (1999). The DCP2 protein is required for mRNA decapping in Saccharomyces cerevisiae and contains a functional MutT motif. EMBO J. 18, 5411–5422.

Dye, B.T., Hao, L., and Ahlquist, P. (2005). High-throughput isolation of Saccharomyces cerevisiae RNA. Biotechniques 38, 868, 870.

Fischer, J., Song, Y.S., Yosef, N., di Iulio, J., Churchman, L.S., and Choder, M. (2020). The yeast exoribonuclease Xrn1 and associated factors modulate RNA polymerase II processivity in 5’ and 3’ gene regions. J. Biol. Chem. 295, 11435–11454.

García-Martínez, J., Pérez-Martínez, M.E., Pérez-Ortín, J.E., and Alepuz, P. (2021a). Recruitment of Xrn1 to stress-induced genes allows efficient transcription by controlling RNA polymerase II backtracking. RNA Biol. 18, 1458–1474.

García-Martínez, J., Medina, D.A., Bellvís, P., Sun, M., Cramer, P., Chávez, S., and Pérez-Ortín, J.E. (2021b). The total mRNA concentration buffering system in yeast is global rather than gene-specific. RNA 27, 1281–1290.

Geisler, S., and Coller, J. (2012). XRN1: A Major 5’ to 3’ Exoribonuclease in Eukaryotic Cells. The Enzymes 31.

Gietz, R.D., Schiestl, R.H., Willems, A.R., and Woods, R.A. (1995). Studies on the transformation of intact yeast cells by the LiAc/SS-DNA/PEG procedure. Yeast 11, 355–360.

Gilbertson, S., Federspiel, J.D., Hartenian, E., Cristea, I.M., and Glaunsinger, B. (2018). Changes in mRNA abundance drive shuttling of RNA binding proteins, linking cytoplasmic RNA degradation to transcription. Elife 7.

Goler-Baron, V., Selitrennik, M., Barkai, O., Haimovich, G., Lotan, R., and Choder, M. (2008). Transcription in the nucleus and mRNA decay in the cytoplasm are coupled processes. Genes Dev. 22, 2022–2027.

Gómez-Herreros, F., Rodríguez-Galán, O., Morillo-Huesca, M., Maya, D., Arista-Romero, M., de la Cruz, J., Chávez, S., and Muñoz-Centeno, M.C. (2013). Balanced production of ribosome components is required for proper G1/S transition in Saccharomyces cerevisiae. J. Biol. Chem. 288, 31689–31700.

Gupta, I., Villanyi, Z., Kassem, S., Hughes, C., Panasenko, O.O., Steinmetz, L.M., and Collart, M.A. (2016). Translational Capacity of a Cell Is Determined during Transcription Elongation via the Ccr4-Not Complex. Cell Rep. 15, 1782–1794.

Gutin, J., Sadeh, A., Rahat, A., Aharoni, A., and Friedman, N. (2015). Condition-specific genetic interaction maps reveal crosstalk between the cAMP/PKA and the HOG MAPK pathways in the activation of the general stress response. Mol. Syst. Biol. 11, 829.

von der Haar, T. (2007). Optimized protein extraction for quantitative proteomics of yeasts. PLoS One 2, e1078.

Haimovich, G., Choder, M., Singer, R.H., and Trcek, T. (2013a). The fate of the messenger is pre-determined: a new model for regulation of gene expression. Biochim. Biophys. Acta 1829, 643–653.

Haimovich, G., Medina, D.A., Causse, S.Z., Garber, M., Millán-Zambrano, G., Barkai, O., Chávez, S., Pérez-Ortín, J.E., Darzacq, X., and Choder, M. (2013b). Gene expression is circular: factors for mRNA degradation also foster mRNA synthesis. Cell 153, 1000–1011.

Harigaya, Y., and Parker, R. (2016). Analysis of the association between codon optimality and mRNA stability in Schizosaccharomyces pombe. BMC Genomics 17, 895.

He, F., Li, X., Spatrick, P., Casillo, R., Dong, S., and Jacobson, A. (2003). Genome-wide analysis of mRNAs regulated by the nonsense-mediated and 5’ to 3’ mRNA decay pathways in yeast. Mol. Cell 12, 1439–1452.

Helenius, K., Yang, Y., Tselykh, T.V., Pessa, H.K.J., Frilander, M.J., and Mäkelä, T.P. (2011). Requirement of TFIIH kinase subunit Mat1 for RNA Pol II C-terminal domain Ser5 phosphorylation, transcription and mRNA turnover. Nucleic Acids Res. 39, 5025–5035.

Henninger, J.E., Oksuz, O., Shrinivas, K., Sagi, I., LeRoy, G., Zheng, M.M., Andrews, J.O., Zamudio, A.V., Lazaris, C., Hannett, N.M., et al. (2021). RNA-Mediated Feedback Control of Transcriptional Condensates. Cell 184, 207–225.e24.

Herzog, V.A., Reichholf, B., Neumann, T., Rescheneder, P., Bhat, P., Burkard, T.R., Wlotzka, W., von Haeseler, A., Zuber, J., and Ameres, S.L. (2017). Thiol-linked alkylation of RNA to assess expression dynamics. Nat. Methods 14, 1198–1204.

Heyer, W.D., Johnson, A.W., Reinhart, U., and Kolodner, R.D. (1995). Regulation and intracellular localization of Saccharomyces cerevisiae strand exchange protein 1 (Sep1/Xrn1/Kem1), a multifunctional exonuclease. Mol. Cell. Biol. 15, 2728–2736.

Hsu, C.L., and Stevens, A. (1993). Yeast cells lacking 5’-->3’ exoribonuclease 1 contain mRNA species that are poly(A) deficient and partially lack the 5’ cap structure. Mol. Cell. Biol. 13, 4826–4835.

Hu, W. (2016). The Interplay between Eukaryotic mRNA Degradation and Translation. Encyclopedia of Cell Biology 346–353.

Huch, S., and Nissan, T. (2014). Interrelations between translation and general mRNA degradation in yeast. Wiley Interdiscip. Rev. RNA 5, 747–763.

Interthal, H., Bellocq, C., Bähler, J., Bashkirov, V.I., Edelstein, S., and Heyer, W.D. (1995). A role of Sep1 (= Kem1, Xrn1) as a microtubule-associated protein in Saccharomyces cerevisiae. EMBO J. 14, 1057–1066.

Jones, C.I., Zabolotskaya, M.V., and Newbury, S.F. (2012). The 5’ → 3’ exoribonuclease XRN1/Pacman and its functions in cellular processes and development. Wiley Interdisciplinary Reviews: RNA 3, 455–468.

Jorgensen, P., Nishikawa, J.L., Breitkreutz, B.-J., and Tyers, M. (2002). Systematic identification of pathways that couple cell growth and division in yeast. Science 297, 395–400.

Jürges, C., Dölken, L., and Erhard, F. (2018). Dissecting newly transcribed and old RNA using GRAND-SLAM. Bioinformatics 34, i218–i226.

Kandasamy, G., Pradhan, A.K., and Palanimurugan, R. (2021). Ccr4-Not complex subunits Ccr4, Caf1, and Not4 are novel proteolysis factors promoting the degradation of ubiquitin-dependent substrates by the 26S proteasome. Biochim. Biophys. Acta Mol. Cell Res. 1868, 119010.

Kemmeren, P., Sameith, K., van de Pasch, L.A.L., Benschop, J.J., Lenstra, T.L., Margaritis, T., O’Duibhir, E., Apweiler, E., van Wageningen, S., Ko, C.W., et al. (2014). Large-scale genetic perturbations reveal regulatory networks and an abundance of gene-specific repressors. Cell 157, 740–752.

Kim, M., Krogan, N.J., Vasiljeva, L., Rando, O.J., Nedea, E., Greenblatt, J.F., and Buratowski, S. (2004). The yeast Rat1 exonuclease promotes transcription termination by RNA polymerase II. Nature 432, 517–522.

Klein-Brill, A., Joseph-Strauss, D., Appleboim, A., and Friedman, N. (2019). Dynamics of Chromatin and Transcription during Transient Depletion of the RSC Chromatin Remodeling Complex. Cell Rep. 26, 279–292.e5.

Koç, A., Wheeler, L.J., Mathews, C.K., and Merrill, G.F. (2004). Hydroxyurea arrests DNA replication by a mechanism that preserves basal dNTP pools. J. Biol. Chem. 279, 223–230.

Kokina, A., Kibilds, J., and Liepins, J. (2014). Adenine auxotrophy--be aware: some effects of adenine auxotrophy in Saccharomyces cerevisiae strain W303-1A. FEMS Yeast Res. 14, 697–707.

Kumar, G.R., Shum, L., and Glaunsinger, B.A. (2011). Importin alpha-mediated nuclear import of cytoplasmic poly(A) binding protein occurs as a direct consequence of cytoplasmic mRNA depletion. Mol. Cell. Biol. 31, 3113–3125.

Labno, A., Tomecki, R., and Dziembowski, A. (2016). Cytoplasmic RNA decay pathways - Enzymes and mechanisms. Biochimica et Biophysica Acta (BBA) - Molecular Cell Research 1863, 3125–3147.

Laribee, R.N., Shibata, Y., Mersman, D.P., Collins, S.R., Kemmeren, P., Roguev, A., Weissman, J.S., Briggs, S.D., Krogan, N.J., and Strahl, B.D. (2007). CCR4/NOT complex associates with the proteasome and regulates histone methylation. Proceedings of the National Academy of Sciences 104, 5836–5841.

Larimer, F.W., and Stevens, A. (1990). Disruption of the gene XRN1, coding for a 5’----3’ exoribonuclease, restricts yeast cell growth. Gene 95, 85–90.

Lu, A.X., Zarin, T., Hsu, I.S., and Moses, A.M. (2019). YeastSpotter: accurate and parameter-free web segmentation for microscopy images of yeast cells. Bioinformatics 35, 4525–4527.

Medina, D.A., Jordán-Pla, A., Millán-Zambrano, G., Chávez, S., Choder, M., and Pérez-Ortín, J.E. (2014). Cytoplasmic 5’-3’ exonuclease Xrn1p is also a genome-wide transcription factor in yeast. Frontiers in Genetics 5.

Metzl-Raz, E., Kafri, M., Yaakov, G., Soifer, I., Gurvich, Y., and Barkai, N. (2017). Principles of cellular resource allocation revealed by condition-dependent proteome profiling.

Morawska, M., and Ulrich, H.D. (2013). An expanded tool kit for the auxin-inducible degron system in budding yeast. Yeast 30, 341–351.

Muhlrad, D., Decker, C.J., and Parker, R. (1994). Deadenylation of the unstable mRNA encoded by the yeast MFA2 gene leads to decapping followed by 5’-->3’ digestion of the transcript. Genes Dev. 8.

Muzzey, D., Gómez-Uribe, C.A., Mettetal, J.T., and van Oudenaarden, A. (2009). A systems-level analysis of perfect adaptation in yeast osmoregulation. Cell 138, 160–171.

Nagarajan, V.K., Jones, C.I., Newbury, S.F., and Green, P.J. (2013). XRN 5’→3’ exoribonucleases: Structure, mechanisms and functions. Biochimica et Biophysica Acta (BBA) - Gene Regulatory Mechanisms 1829, 590–603.

Newbury, S. (2004). The 5’-3’ exoribonuclease xrn-1 is essential for ventral epithelial enclosure during C.elegans embryogenesis. RNA 10, 59–65.

Nishimura, K., Fukagawa, T., Takisawa, H., Kakimoto, T., and Kanemaki, M. (2009). An auxin-based degron system for the rapid depletion of proteins in nonplant cells. Nat. Methods 6, 917–922.

Padovan-Merhar, O., Nair, G.P., Biaesch, A.G., Mayer, A., Scarfone, S., Foley, S.W., Wu, A.R., Churchman, L.S., Singh, A., and Raj, A. (2015). Single mammalian cells compensate for differences in cellular volume and DNA copy number through independent global transcriptional mechanisms. Mol.Cell 58, 339–352.

Panasenko, O.O., and Collart, M.A. (2011). Not4 E3 ligase contributes to proteasome assembly and functional integrity in part through Ecm29. Mol. Cell. Biol. 31, 1610–1623.

Panasenko, O.O., Somasekharan, S.P., Villanyi, Z., Zagatti, M., Bezrukov, F., Rashpa, R., Cornut, J., Iqbal, J., Longis, M., Carl, S.H., et al. (2019). Co-translational assembly of proteasome subunits in NOT1-containing assemblysomes. Nat. Struct. Mol. Biol. 26, 110–120.

Parker, R. (2012). RNA Degradation in Saccharomyces cerevisae. Genetics 191, 671–702.

Picelli, S., Björklund, A.K., Reinius, B., Sagasser, S., Winberg, G., and Sandberg, R. (2014). Tn5 transposase and tagmentation procedures for massively scaled sequencing projects. Genome Res. 24, 2033–2040.

Plaschka, C., Larivière, L., Wenzeck, L., Seizl, M., Hemann, M., Tegunov, D., Petrotchenko, E.V., Borchers, C.H., Baumeister, W., Herzog, F., et al. (2015). Architecture of the RNA polymerase II–Mediator core initiation complex. Nature 518, 376–380.

Rahman, S., and Zenklusen, D. (2013). Single-molecule resolution fluorescent in situ hybridization(smFISH) in the yeast S. cerevisiae. Methods Mol. Biol. 1042, 33–46.

Raj, A., van den Bogaard, P., Rifkin, S.A., van Oudenaarden, A., and Tyagi, S. (2008). Imaging individual mRNA molecules using multiple singly labeled probes. Nat. Methods 5, 877–879.

Rodríguez-Molina, J.B., Tseng, S.C., Simonett, S.P., Taunton, J., and Ansari, A.Z. (2016). Engineered Covalent Inactivation of TFIIH-Kinase Reveals an Elongation Checkpoint and Results in Widespread mRNA Stabilization. Mol. Cell 63, 433–444.

Santos, A., Wernersson, R., and Jensen, L.J. (2015). Cyclebase 3.0: a multi-organism database on cell-cycle regulation and phenotypes. Nucleic Acids Res. 43, D1140–D1144.

Schmid, M., and Jensen, T.H. (2018). Controlling nuclear RNA levels. Nature Reviews Genetics 19, 518–529.

Schmid, M., Olszewski, P., Pelechano, V., Gupta, I., Steinmetz, L.M., and Jensen, T.H. (2015). The Nuclear PolyA-Binding Protein Nab2p Is Essential for mRNA Production. Cell Rep. 12, 128–139.

Schulz, D., Pirkl, N., Lehmann, E., and Cramer, P. (2014). Rpb4 subunit functions mainly in mRNA synthesis by RNA polymerase II. J. Biol. Chem. 289, 17446–17452.

Shalem, O., Dahan, O., Levo, M., Martinez, M.R., Furman, I., Segal, E., and Pilpel, Y. (2008). Transient transcriptional responses to stress are generated by opposing effects of mRNA production and degradation. Mol. Syst. Biol. 4, 223.

Shalem, O., Groisman, B., Choder, M., Dahan, O., and Pilpel, Y. (2011). Transcriptome kinetics is governed by a genome-wide coupling of mRNA production and degradation: a role for RNA Pol II. PLoS Genet. 7, e1002273.

Sheth, U., and Parker, R. (2003). Decapping and decay of messenger RNA occur in cytoplasmic processing bodies. Science 300, 805–808.

Slobodin, B., Bahat, A., Sehrawat, U., Becker-Herman, S., Zuckerman, B., Weiss, A.N., Han, R., Elkon, R., Agami, R., Ulitsky, I., et al. (2020). Transcription Dynamics Regulate Poly(A) Tails and Expression of the RNA Degradation Machinery to Balance mRNA Levels. Mol. Cell 78, 434–444.e5.

Stevens, A. (1978). An exoribonuclease from saccharomyces cerevisiae: Effect of modifications of 5’ endgroups on the hydrolysis of substrates to 5’ mononucleotides. Biochemical and Biophysical Research Communications 81, 656–661.

Stevenson, K., McVey, A.F., Clark, I.B.N., Swain, P.S., and Pilizota, T. (2016). General calibration of microbial growth in microplate readers. Sci. Rep. 6, 38828.

Stovner, E.B., and Sætrom, P. (2020). PyRanges: efficient comparison of genomic intervals in Python. Bioinformatics 36, 918–919.

Sun, M., Schwalb, B., Schulz, D., Pirkl, N., Etzold, S., Larivière, L., Maier, K.C., Seizl, M., Tresch, A., and Cramer, P. (2012). Comparative dynamic transcriptome analysis (cDTA) reveals mutual feedback between mRNA synthesis and degradation. Genome Res. 22, 1350–1359.

Sun, M., Schwalb, B., Pirkl, N., Maier, K.C., Schenk, A., Failmezger, H., Tresch, A., and Cramer, P. (2013). Global analysis of eukaryotic mRNA degradation reveals Xrn1-dependent buffering of transcript levels. Mol. Cell 52, 52–62.

Trcek, T., Larson, D.R., Moldón, A., Query, C.C., and Singer, R.H. (2011). Single-molecule mRNA decay measurements reveal promoter- regulated mRNA stability in yeast. Cell 147, 1484–1497.

Tucker, M., Valencia-Sanchez, M.A., Staples, R.R., Chen, J., Denis, C.L., and Parker, R. (2001). The transcription factor associated Ccr4 and Caf1 proteins are components of the major cytoplasmic mRNA deadenylase in Saccharomyces cerevisiae. Cell 104, 377–386.

Voichek, Y., Bar-Ziv, R., and Barkai, N. (2016). Expression homeostasis during DNA replication. Science 351, 1087–1090.

Wada, T., and Becskei, A. (2017). Impact of Methods on the Measurement of mRNA Turnover. Int. J. Mol. Sci. 18, 2723.

Warfield, L., Ramachandran, S., Baptista, T., Devys, D., Tora, L., and Hahn, S. (2017). Transcription of Nearly All Yeast RNA Polymerase II-Transcribed Genes Is Dependent on Transcription Factor TFIID. Molecular Cell 68, 118–129.e5.

Wasmuth, E.V., Januszyk, K., and Lima, C.D. (2014). Structure of an Rrp6–RNA exosome complex bound to poly(A) RNA. Nature 511, 435–439.

Weiner, A., Hsieh, T.-H.S., Appleboim, A., Chen, H.V., Rahat, A., Amit, I., Rando, O.J., and Friedman, N. (2015). High-resolution chromatin dynamics during a yeast stress response. Mol. Cell 58, 371–386.

Weiße, A.Y., Oyarzún, D.A., Danos, V., and Swain, P.S. (2015). Mechanistic links between cellular trade-offs, gene expression, and growth. Proc. Natl. Acad. Sci. U. S. A. 112, E1038–E1047.

Wiener, D., Antebi, Y., and Schwartz, S. (2021). Decoupling of degradation from deadenylation reshapes poly(A) tail length in yeast meiosis. Nat. Struct. Mol. Biol. 28, 1038–1049.

Yang, E., van Nimwegen, E., Zavolan, M., Rajewsky, N., Schroeder, M., Magnasco, M., and Darnell, J.E., Jr (2003). Decay rates of human mRNAs: correlation with functional characteristics and sequence attributes. Genome Res. 13, 1863–1872.

Young, E.T., Zhang, C., Shokat, K.M., Parua, P.K., and Braun, K.A. (2012). The AMP-activated protein kinase Snf1 regulates transcription factor binding, RNA polymerase II activity, and mRNA stability of glucose-repressed genes in Saccharomyces cerevisiae. J. Biol. Chem. 287, 29021–29034.

Zid, B.M., and O’Shea, E.K. (2014). Promoter sequences direct cytoplasmic localization and translation of mRNAs during starvation in yeast. Nature 514, 117–121.

