## supplementary text and figures for "Transcription feedback dynamics in the wake of cytoplasmic degradation shutdown"

#### Supplementary Material


**Supplementary Figures** **2**
- Figure S1. cDTA-seq validation and analysis 2
- Figure S2. Xrn knockouts analyses 4
- Figure S3. transient mRNA accumulation upon Xrn1 depletion 6
- Figure S4. Excluding non-transcriptional explanations to the reduction in RNA levels 8
- Figure S5: Cell cycle is linked to the transcription adaptation response 10
- Figure S6: The transcription adaptation response along the 5’-3’ branch 12

**Supplementary Tables** **14**
- Table S1. Yeast strains used in this study 14
- Table S2. Oligonucleotides used in this study 15
- Table S3. FISH probes 16

**Supplementary Notes** **17**
- Half-life estimation from 4tU labeling data 17
- First order model predictions when degradation is reduced but production is not 19
- Additional analyses 21
    - Not1 sensitive transcripts 21

**Supplementary References** **22**

### Supplementary Figures

#### Figure S1. cDTA-seq validation and analysis

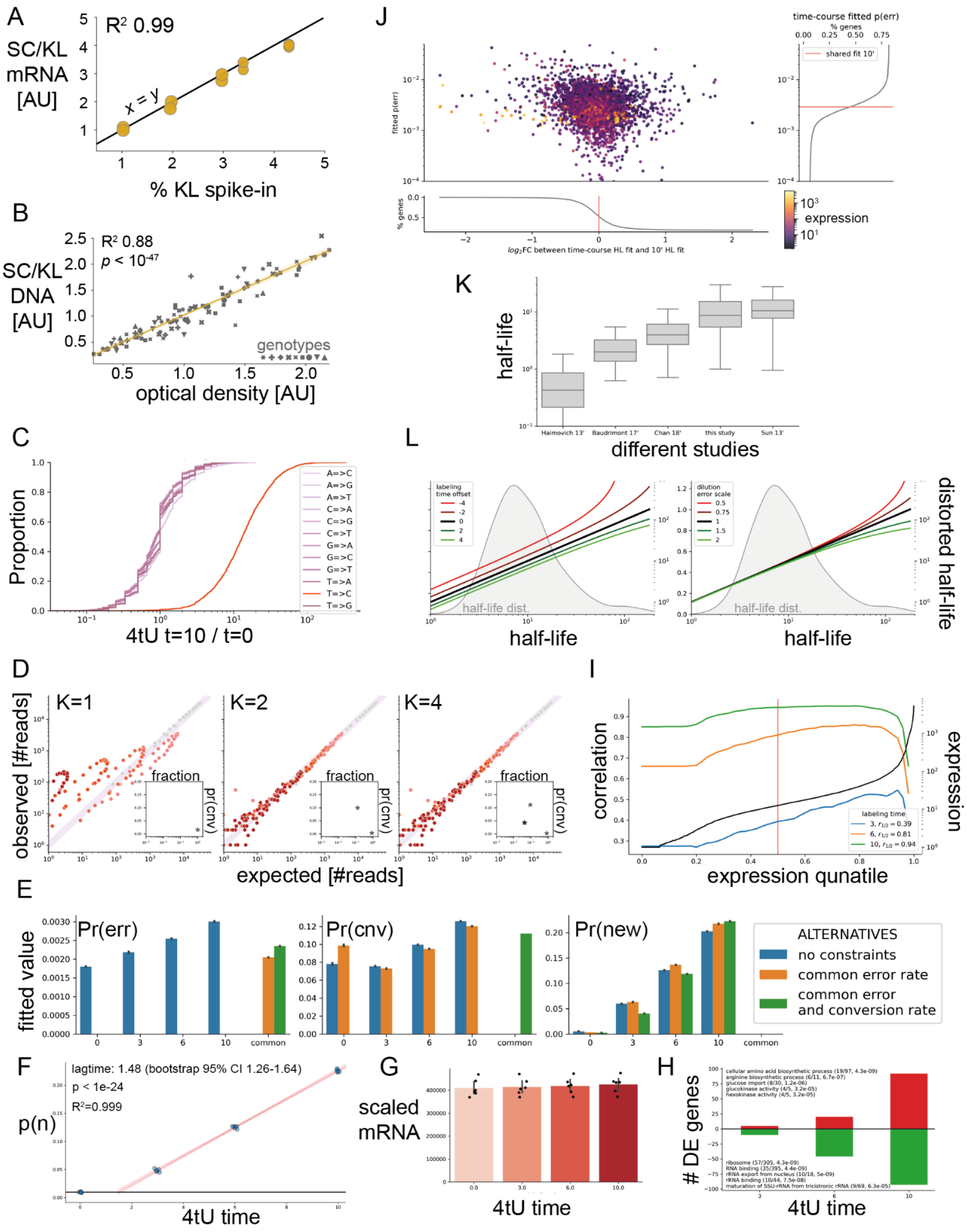

**Supplementary Figure 1 (related to Figure 1): cDTA-seq validation and analysis**

**A) Spike-in read ratio is linear.** A SC sample was split and spiked in with varying amounts of KL cells (1-5%, x-axis). The ratio of mRNA reads from SC and KL (y-axis) is linear in this range (N=2, $R^2$=0.98, black line is x=y).

**B) OD is a good proxy for the number of cells.** Each marker is one of 93 samples with various genotypes (various markers) in various conditions (methods). Samples' optical density (OD, x-axis) was measured, they were spiked in with KL cells, DNA was extracted and the SC/KL DNA ratio (y-axis) is plotted against their measured OD ($R^2$=0.88, $p < 10^{-47}$, yellow line is linear fit, shaded with 95% CI).

**C) Only T→C observations are changed after 4tU labeling.** Observed deviations from the reference were counted per transcript. The change in this count after 10 minutes of 4tU labeling is plotted as log fold change (x-axis) cumulative distribution across all transcripts (y-axis).

**D) BMM with more than 2 components doesn't perform better.** The binomial mixture model was fitted to the data (after 10' of labeling) with a varying number of mixture components (shown are 1, 2, and 4). Given a fitted model and the distribution of observed Ts per read, one can plot the expected number of reads per combination of Ts and converted bases (y-axis) vs. the actual number of observed reads per each combination (x-axis). Insets show the fitted parameters. Each component is represented in the space of the fraction of the population (x-axis) and the T to C conversion rate it was fitted with (methods).

**E) Alternative parameter sharing schemes show dominant change is due to $p_n$.** 2-BMM was fitted to the 4tU time course data without any shared parameters (i.e. full model per sample, blue), with a shared error rate ("ε", orange), or with shared error rate and a shared conversion rate ("ξ", green). In all cases, the pr(new) ("$p_n$") fitted parameter (right) correlates with 4tU labeling time (x-axis). N=6.

**F) 4tU lag-time estimation.** Fitting a linear model to the time-course data (confidence interval from data resampling) predicts that the labeling lag time is ~1.5 minutes.

**G) No change to global mRNA.** Spike-in scaled estimates of mRNA levels in samples along the 4tU time course show no changes to overall mRNA levels.

**H) Differentially expressed gene increase with 4tU (<200 genes at 10').** DESeq2 differential expressed genes compared to t=0 along the 4tU time course with 6 replicates. After ~10 minutes of labeling ~ 180 transcripts are up (red) or down (green) regulated compared to t=0. Enriched GO annotations in each set are denoted in text.

**I) Correlation between whole-time-course and single-sample-estimate is high between 6'-10' labeling.** The half life of each transcript was estimated with the full time course or with a single time point (3',6',10'), and the correlation between these estimates (y-axis) was plotted as a function of expression cutoff (x-axis). For example at x=0.2, the correlation was performed only considering the top 80% of expressed transcripts. Black line denotes the expression levels [AU, log scale].

**J) p(err) is not correlated to the difference between whole-time-course and single-sample-estimate, nor to expression levels.** Each transcript 4tU time course data was fitted a model allowing for a transcript-specific error rate ("ε"), rather than a global one. Fitted values (y-axis, CDF with median in red on the right) range from 0.001 to 0.007 which are at least an order of magnitude lower than the fitted conversion rate ("ξ", ~0.11). Differences between transcripts are not correlated to half-life error estimations (x-axis, log fold change of half life estimate based on full time course vs. half life based on a single 10 min time point). Differences are also not correlated to expression levels (log scale, color coded).

**K) Global differences in half-life distributions between studies.** The distributions of half life estimates (in minutes, y-axis) are box-plotted for 5 different studies to highlight the different estimates scales.

**L) Half-life estimation is robust to global parameter estimations.** The theoretic halflife estimation (right y-axis) is compared to the actual half life (x-axis), assuming different errors in global parameter estimations. For example, if cell growth is actually slower by a factor of two compared to the rate used when estimating half life, the distortion is given by the bright green line on the right panel. Background smoothed histogram depicts the distribution of half lives across the genome, guiding the eye to where most transcripts reside.

### Figure S2. Xrn knockouts analyses

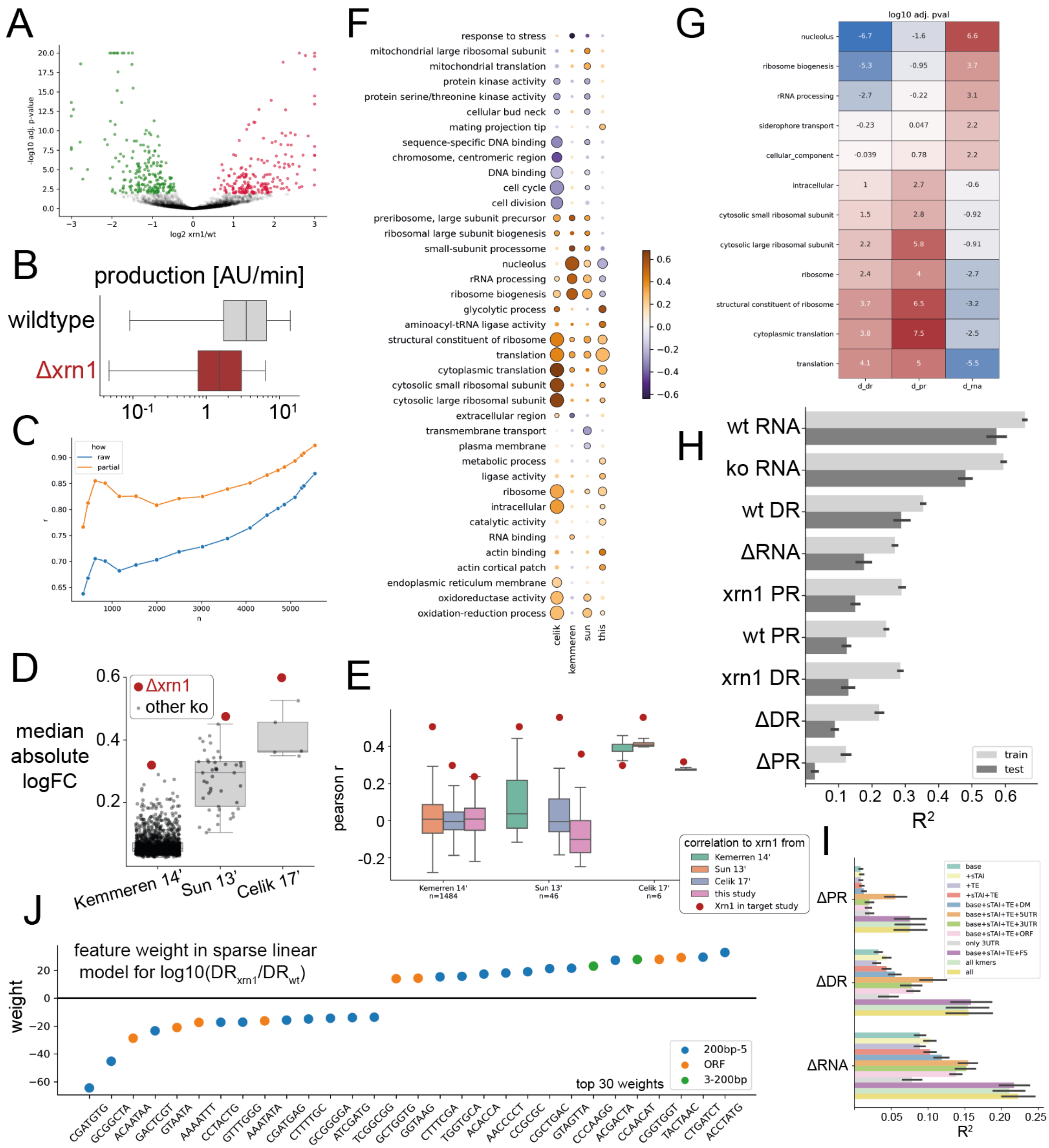

**Supplementary Figure 2 (related to Figure 2): Xrn1 knockouts analyses**

**A) Xrn1 knockout differential expression.** Comparing two wildtype biological replicates and two biological replicates of two different clones without Xrn1, ~400 transcripts were significantly changed (adjusted $p < 10^{-2}$ (Love et al., 2014)).

**B) Inferred production rates are significantly reduced in Xrn1 knockout.** Transcript production rate distributions, as inferred from mRNA levels and estimated degradation rates.

**C) Correlation between change to production and degradation rate.** Conditioned on genes with expression > threshold (n - number of genes above the threshold). The orange curve is the partial correlation(Vallat, 2018), conditioned on mRNA levels.

**D) An extreme transcriptomic signature for Xrn1 knockout.** In three different knockout studies in yeast (x-axis), Xrn1 knockout (red dots) exerts the largest effect on the measured transcriptome as measured by the median over the absolute log2 fold change relative to wildtype strain (y-axis).

**E) Correlations between studies.** Correlating an Xrn1 KO from each "source" study (box color) to all KO strains in other studies ("target study", x-axis) generates a distribution of correlations (boxes). For example, correlating Xrn1 from Sun 13' with all KOs in Kemerren 14' generates the left-most orange box plot. Xrn1 in the target study is highlighted as a red dot, demonstrating that while correlation between studies is low, Xrn1 is still the most significant correlation in most comparisons.

**F) Enriched functional sets knockout studies are incongruent.** Enriched sets in up- or down-regulated genes in each study (15 top per study). Circle size is proportional to negative log BH-adjusted p-value (kolmogorov-smirnov test of set relative to overall distribution). Color denotes the signed KS statistic (orange - higher than average, purple lower than average).

**G) Extreme gene sets in Xrn1 KO changes.** All GO annotations (rows) were tested for having extreme distributions in their degradation rates, their mRNA levels, or their production rates (columns) using a kolmogorov-smirnov test. BH adjusted q-values are color-coded such that below average changes are blue, and above average changes are red (colorbar). Numbers indicate log10 q-value. Only significant sets are shown (rows).

**H) Explaining observed RNA features.** Trying to explain various observed RNA measurements (y-axis) with a sparse linear model (LASSO) using various features and sequences (see supplementary note on modeling ΔXrn1 changes). X-axis is the % explained variance by the model prediction ($R^2$), on the training data (light grey, 90% of genes in 10 random samples), or the remaining test data (dark grey, 10% of genes).

**I) Sequence features are sufficient for model performance.** Plotting model performance (x-axis, $R^2$) Using different subsets of features. K-mer counts from ORF, 5', and 3' areas are sufficient for model performance (see supplementary note on modeling ΔXrn1 changes).

**J) K-mer examples.** K-mers (x-axis) used by the model for explaining changes to degradation rates, ordered by their relative weight (y-axis), and colored by their class (ORF/5'/3'). Showing only top 30 out of ~200 LASSO-selected k-mers in this case (degradation rate changes).

### Figure S3. transient mRNA accumulation upon Xrn1 depletion

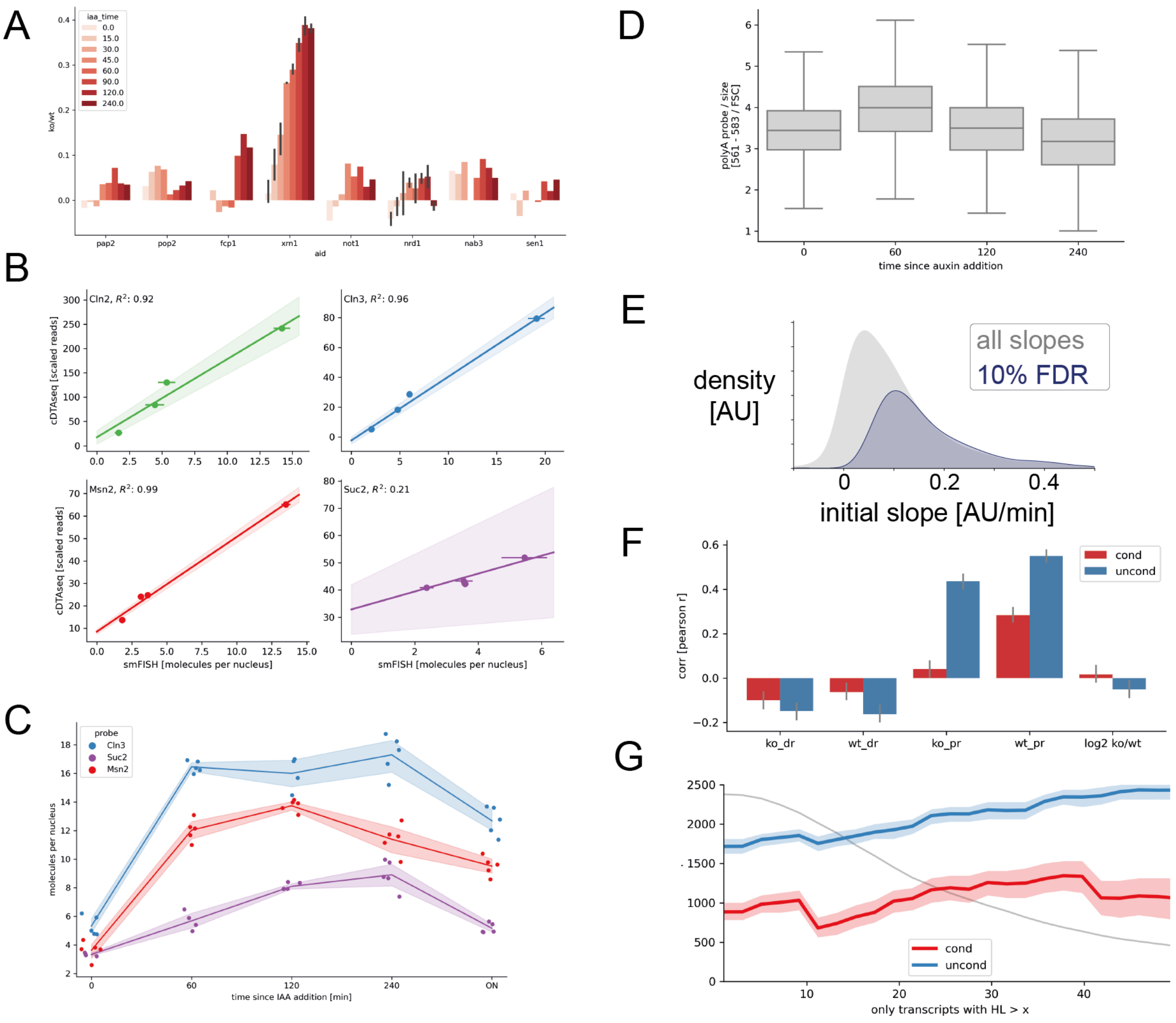


**Supplementary Figure 3 (related to Figure 3): transient mRNA accumulation upon Xrn1 depletion**

**A) Correlation of Xrn1 depletion to Xrn1 KO.** log FC change from corresponding wt samples of multiple measurements following auxin addition of multiple degron strains were correlated to the Xrn1 knockout/wt log fold change. The correlation between the Xrn1-AID strain and the Xrn1 KO strain increases with time, i.e. these expression profiles become similar to that of the KO strain.

**B) smFISH cDTA-seq comparison.** smFISH measurements (x-axis) and cDTA-seq measurements (y-axis) are highly correlated. Points are one of four samples: Spt6-AID or Xrn1-AID with mock or auxin treatment for 60' minutes.

**C) Slower RNA decrease in smFISH time course.** Auxin time course (x-axis, last point is overnight) smFISH spots/nucleus (y-axis) for three probes shows the increase and slower decrease in total RNA.

**D) polyA signal follows the cDTA-seq dynamics.** polyA fluorescent probes quantified by FACS and normalized to forward scatter (y-axis). Distribution of 10k cells per time point shows an increase and decrease by 240 minutes, consistent with cDTA-seq data.

**E) Slopes are positive and linear.** The distribution of fitted initial slopes (x-axis, light grey) demonstrates that virtually all transcripts increase following Xrn1 depletion. We select transcripts that can be reasonably fitted with a linear fit (linear fit p-value, false discovery rate of 10%) for the following analysis. Colored dots indicate the slope (and FDR status) of the transcripts shown in (G).

**F) Initial slope correlates to production rate, also after accounting for mRNA levels.** Correlation (y-axis) between transcript slope to different transcript rates and measures (x-axis) was calculated with (red) or without (blue) conditioning on mRNA levels.

**G) Initial slope correlation to production rate increases with transcript stability.** As in (E), only examining subsets of transcripts with increasing stability (x-axis denotes the half-life threshold). Grey line indicates the number of transcripts (y-axis) used as threshold increases.

### Figure S4. Excluding non-transcriptional explanations to the reduction in RNA levels

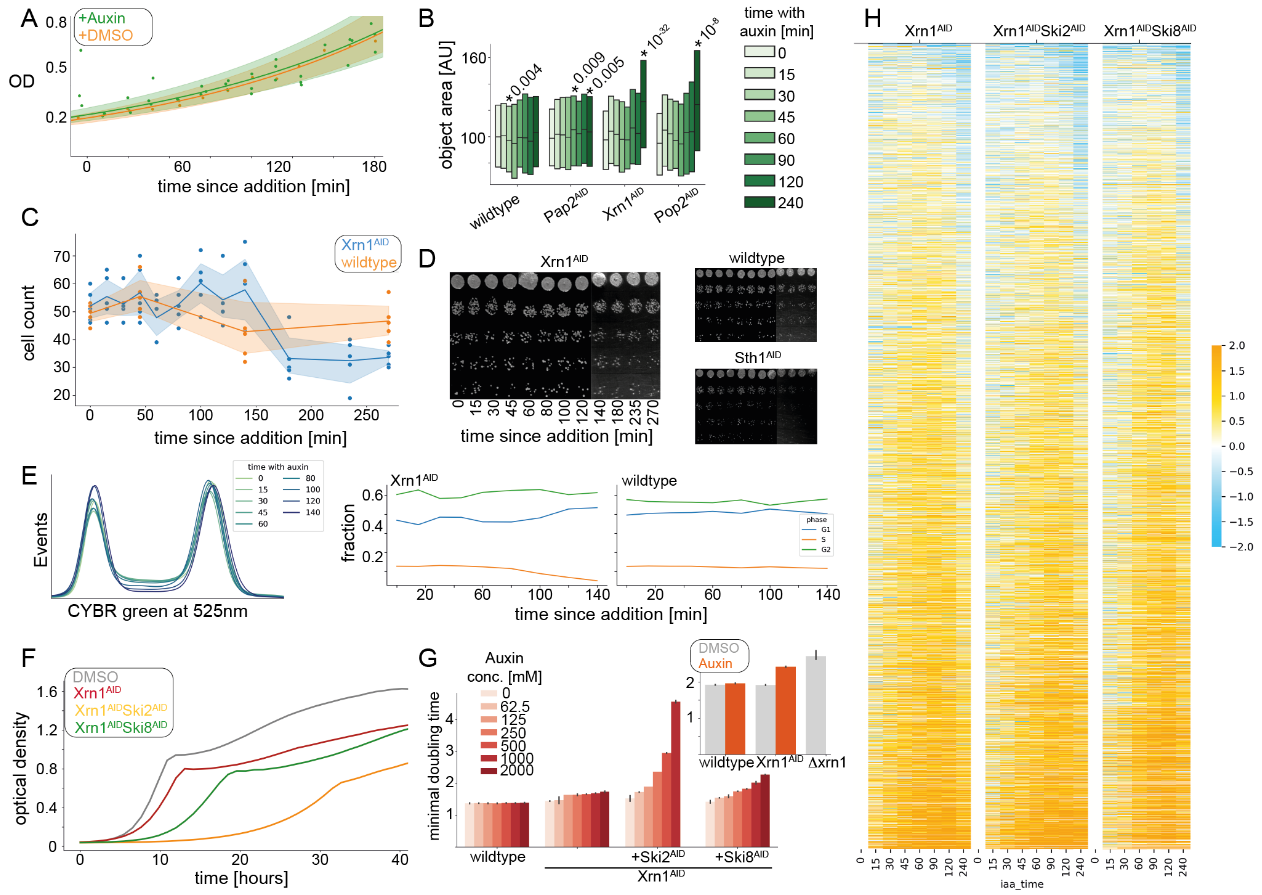

**Supplementary Figure 4 (related to Figure 4): Excluding non-transcriptional explanations to reduction in RNA levels**

**A) No immediate changes to OD or growth.** OD measurements (y-axis) for samples grown on plates for 2 hours (x-axis starts at 120 minutes), and then supplemented with auxin (green) or not (orange). No significant changes in growth are observed within 3 hours.

**B) Significant size changes observed only after 4 hours.** Several strains were imaged (supplementary methods), objects were segmented (Lu et al., 2019), and their area was quantified (y-axis). Each box group is a strain, colors indicate time since auxin addition. Each time-course was normalized to the median area at times (0,15). Significant t-test p-values are noted on the diagram. Xrn1 and Pop2 knockouts are known to have increased size (Jonas et al., 2018; Jorgensen, 2002; Zhang et al., 2002).

**C) No significant changes to cell counts within 2 hours.** Cells were fixed in formaldehyde, vortexed, blinded, and manually counted on a hemocytometer (y-axis) along a time course with auxin (x-axis). A clear reduction in Xrn1$^{AID}$ cell counts is observed only after ~3 hours.

**D) Colony forming units show little change along an auxin time course.** Cells were exposed to auxin for various durations (x-axis), serially diluted and plated on YPD plates (no-auxin). A slight reduction is observed in the last three time points. As a control - no changes are observed in the wildtype strain, and a significant reduction is observed in an Sth1$^{AID}$ strain (as previously observed (Klein-Brill et al. 2019)).

**E) No gross changes to cell cycle proportions in the first 2 hours following auxin addition.** DNA staining (supplementary methods) of cells following auxin addition shows no gross changes to DNA distribution in an Xrn1$^{AID}$ population (left histograms). Fitted proportions of cells (supplementary methods) are fixed in a wildtype control (right) but potentially show a slight decrease in S-phase cells in Xrn1$^{AID}$ after ~80 minutes (left).

**F) Xrn1-AID, Xrn1-Ski2-AID, and Xrn1-Ski8-AID growth curves.** Growth curves (y-axis is OD, x-axis is time in hours) for an isogenic strain (wilftype, grey), Xrn1-AID (red), double AID tag for Xrn1 and Ski8 (green), and double AID tag for Xrn1 and Ski2 (yellow) with final auxin concentration of 1000 µM.

**G) Effects of different auxin concentration on growth.** Maximal doubling time (y-axis) recorded in the growth curves (examples in (D)) of the indicated strains (x-axis, same as in (D)) with difference concentration of auxin (colors). Inset: maximal doubling time (y-axis) for wildtype, Xrn1-AID, and Xrn1 knockout with (orange) and without (grey) auxin (2.2 mM).

**H) No differences in RNA profile following auxin in double-tagged strains.** RNA profile in time (x-axis) as log2 fold change relative to t=0 (color-coded) per transcript (rows) in the Xrn1-AID strain (left), and the double tagged strains (+Ski2-AID, +Ski8-AID). Same as Figure 3D.

### Figure S5: Cell cycle is linked to the transcription adaptation response

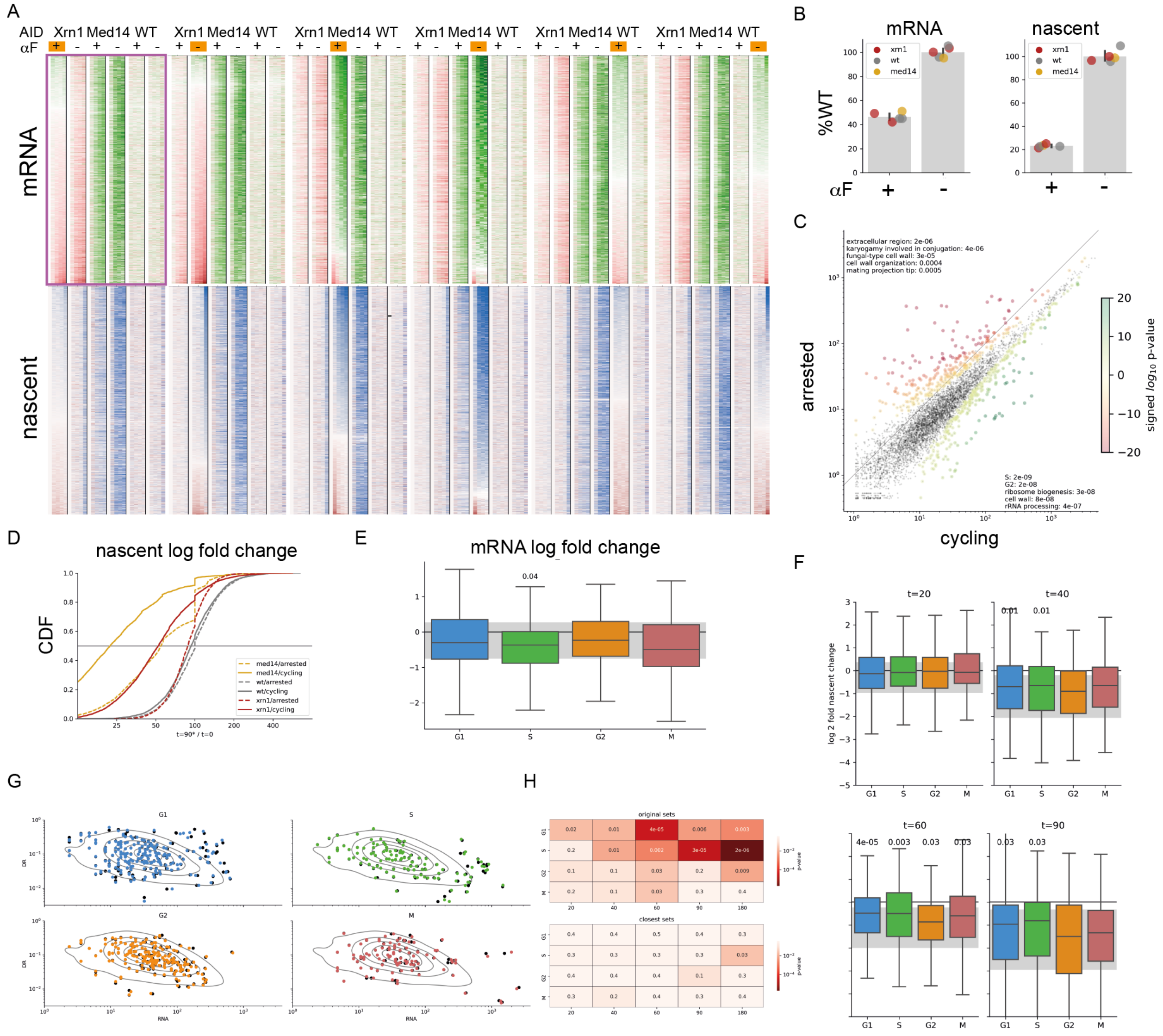

**Supplementary Figure 5 (related to Figure 5): Cell cycle is linked to the transcription adaptation response**

**A) Cell cycle experiment data sorted by every sample.** Each heatmap entry is the change in mRNA (top, red/green) or in nascent mRNA (bottom, red/blue) relative to t=0. Rows are transcripts, columns are sorted by strains and then by time since auxin addition (20/40/60/90 minutes). Each heatmap is sorted by the signed maximal change in one of the samples (indicated in orange). For example the top left heatmap (purple box) is sorted by the mRNA change in the Xrn-AID strain when it was arrested (+αF, orange).

**B) Basal state of arrested cells.** Total and nascent mRNA are significantly lowered in the arrested cells (+) compared to cycling cells (-), prior to auxin addition . Colors indicate different AID strains. Bars indicate average value and error bars indicate the SEM (N=5). Mean decreases to 46.5% of cycling cells total mRNA (t-test $p < 10^{-7}$), and 23.1 % of nascent mRNA (t-test $p < 10^{-8}$).

**C) Differential expression highlights the expected cell-cycle arrest signatures in mRNA profiles.** Average expression per transcript (dots) over all strains and repeats in cycling (x-axis) or arrested (y-axis) cells. Data was analyzed with DESeq2 (https://doi.org/doi:10.18129/B9.bioc.DESeq2) analysis, and the adjusted p-values were used to color genes that are differentially expressed (colorbar is log-scaled, red - significant in arrested, green - significant in cycling). Upper left text lists GO annotations enriched in the up-regulated genes upon arrest, bottom right lists annotations enriched in down-regulated genes.

**D) It is possible to measure significant nascent mRNA reduction in arrested cells.** Cumulative distribution functions (CDFs) for change in nascent mRNA of the three strains (colored as in (A) and (B)) after 90 minutes of auxin. Each strain has a cycling sample (solid line) and arrested sample (dashed line). As in Figure 5F, since the cycling wildtype 90' sample had too few reads, the 60' minute sample was used in this plot. The plot show a dramatic decrease in nascent mRNA upon Med14 depletion from arrested cells (yellow dashed line).

**E) No major outliers in cell-cycle gene sets in Xrn1 knockout.** Log fold change (y-axis) between Xrn1 knockout and wildtype in mRNA levels, grouped by cell cycle sets (colored boxes). Significant adjusted p-values (kolmogorov-smirnov, compared to all genes in grey) denoted as text above respective box.

**F) Sth1 shows a reversed G1/S signature.** Repeating the analysis from Figure 5H on the Sth1 depletion time course shows that the G1/S genes are higher than the non-cycling genes. Significant adjusted p-values denoted as text above respective box. Time point indicated above each figure.

**G) Selection of matched genes to cell cycle sets.** Nearest neighbour procedure was used to get a very close distribution of genes in the mRNA (x-axis) vs. degradation rate (y-axis) space. Colored dots denote the named set, black dots are the matched set.

**H) Matched sets demonstrate that cell-cycle signature is not due to degradation rate or mRNA levels.** Original sets (top), or matched sets (I) were subjected to the same analysis as in (H) - no significant differences between matched sets and all other genes (bottom heatmap - no significant p-values).

### Figure S6: The transcription adaptation response along the 5’-3’ branch

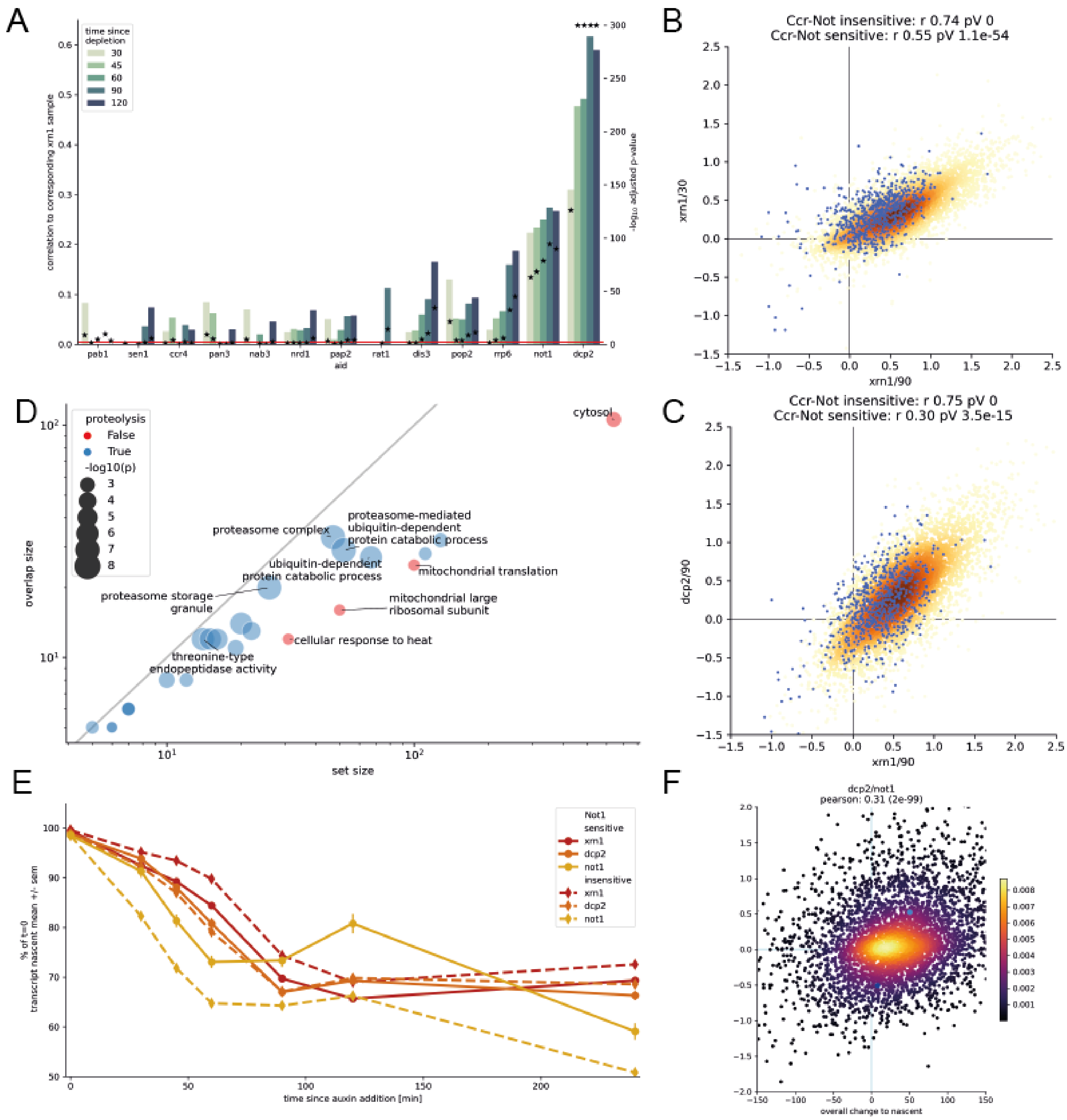


**Supplementary Figure 6 (related to Figure 6): The transcription adaptation response along the 5’-3’ branch**

**A) Correlations to Xrn1.** Correlations along the depletion time course (color legend) of various proteins (x-axis) to Xrn1 (in corresponding time points). Left y-axis denotes the Pearson correlation value(bars), right y-axis denotes the -log10 p-value of the correlation (black stars). Red horizontal line is the adjusted threshold for significant p-values.

**B-C) RNA log2 fold change between various samples.** Scatters exemplifying correlations shown in panel A and figure 6B. Same as Figure 6D. The set of Not1 sensitive transcripts (blue dots) is only exceptional in (D), but maintains overall trend, suggesting an additive effect of 5'-3' interference response and Not1-specific response.

**D) Not1-sensitive transcripts are enriched in proteolysis annotations.** The set of Not1-sensitive genes was tested for significant overlaps with GO annotations (hypergeometric tests). Most enriched sets are proteolysis related (blue). Log10 p-values are proportional to set marker size. Top 10 sets are named on plot.

**E) Nascent mRNA dynamics in Not1 sensitive transcripts.** Same as Figure 6E (dashed lines here correspond to solid lines in Figure 6E), but also showing the dynamics of Not1-sensitive transcripts (solid lines). Due to the selection process, an opposite difference between Xrn1 and Not1 is generated (methods), but overall the trends observed throughout the nascent transcriptome are maintained also in this Not1-sensitive set.

**F) Nascent mRNA differences explain mRNA differences between strains.** Same as Figure 6H, but comparing Not1 to Dcp2.

### Supplementary Tables

#### Table S1. Yeast strains used in this study

All strains in this study were derived from a gift from Ulrich ("nf164") with the following genotype (DF, mat-a): his3-Δ200, leu2-3,2-112, lys2-801, trp1-1(am), URA3::TIR-9Myc.

| NF number | Name | Parental strain | Genotype |
|---|---|---|---|
| nf227 | XRN1-IAA*-MYC | nf164 | XRN1-44AID9Myc::natNT |
| nf198 | STH1-IAA*-FLAG | nf164 | STH1-44AID9Flag::hphNT |
| nf189 | SPT6-IAA*-FLAG | nf164 | SPT6-44AID9Flag::hphNT |
| aa40 | dcp2-degM | nf164 | DCP2-44AID9Myc::natNT |
| aa44 | rat1-degF | nf164 | RAT1-44AID9Flag::hphNT |
| aa46 | med14-degF | nf164 | MED14-44AID9Flag::hphNT |
| aa47 | med14-degM | nf164 | MED14-44AID9Myc::natNT |
| aa82 | Pop2-AID | nf164 | POP2-44AID9Myc::natNT |
| aa85 | fcp1-AID | nf164 | FCP1-44AID9Flag::hphNT |
| aa72 | NatΔXrn1 | nf164 | Xrn1::natNT |
| aa73 | HygΔXrn1 | nf164 | Xrn1::hphNT |
| aa106 | Xrn1-Flag-HygMX | nf164 | Xrn1-44AID9Flag::hphNT |
| aa88 | Xrn1-Ski2-AID | nf227 | XRN1-44AID9Myc::natNT, SKI2-44AID9Myc::kanMX |
| aa90 | Xrn1-Ski8-AID | nf227 | XRN1-44AID9Myc::natNT, SKI8-44AID9Myc::kanMX |
| aa100 | Pap2-AID | nf164 | PAP2-44AID9Myc::natNT |
| aa92 | Nrd1-Flag-HygMX | nf164 | NRD1-44AID9Flag::hphNT |
| aa93 | Nrd1-Myc-NatMX | nf164 | NRD1-44AID9Myc::natNT |
| aa98 | Sen1-Flag-HygMX | nf164 | SEN1-44AID9Flag::hphNT |
| aa97 | Nab3-Myc-NatMX | nf164 | NAB3-44AID9Myc::natNT |
| aa113 | Dis3-Myc-NatMX | nf164 | DIS3-44AID9Myc::natNT |
| aa114 | Rrp6-Myc-NatMX | nf164 | RRP6-44AID9Myc::natNT |
| aa118 | Ccr4-Myc-NatMX | nf164 | CCR4-44AID9Myc::natNT |
| aa91 | Not1-Myc-NatMX | nf164 | NOT1-44AID9Myc::natNT |
| aa116 | Pab1-Myc-NatMX | nf164 | PAB1-44AID9Myc::natNT |
| aa119 | Pan3-Myc-NatMX | nf164 | PAN3-44AID9Myc::natNT |

### Table S2. Oligonucleotides used in this study

The following primer pairs were used to AID-tag the various strain shown in table S1:

| Name | Sequence |
|---|---|
| xrn1-deg-F | CAATGCTGCTGACCGTGATAATAAAAAAGACGAATCTACTcgtacgctgcaggtcgac |
| xrn1-deg-R | TAAAGTAACCTCGAATATACTTCGTTTTTAGTCGTATGTTatcgatgaattcgagctcg |
| STH1-deg-F | AAATGAGTTTACTGATGAATGGTTCAAGGAACACTCTTCGcgtacgctgcaggtcgac |
| STH1-deg-R | ATATAGTCGTAAAAAAAAAAAACATGTGGTGATGAAAACGatcgatgaattcgagctcg |
| SPT6-deg-F | AAAATCTAACAGTAGTAAGAATAGAATGAACAACTACCGTcgtacgctgcaggtcgac |
| SPT6-deg-R | ATAATAAAATTAATAATAACAATGGACACTACATACGCATatcgatgaattcgagctcg |
| dcp2-f-deg | TTCAGGGTCTAATGAATTATTAAGCATTTTGCATAGGAAGcgtacgctgcaggtcgac |
| dcp2-r-deg | CATTTACAGTGTGTCTATAAAACGTATAACACTTATTCTTatcgatgaattcgagctcg |
| xrn1_ko_F | ACTTGTAACAACAGCAGCAACAAATATATATCAGTACGGTcgtacgctgcaggtcgac |
| rat1-f-deg | CAAGCAAAGTCGGTATGACAATTCAAGAGCAAATAGGCGTcgtacgctgcaggtcgac |
| rat1-r-deg | AACCTAAATTTACCATAAAATAAAATGCGCACGAGTAGTTatcgatgaattcgagctcg |
| med14-f-deg | CCATAATATCCTCAAAGTGGACTCGAACTCAAGTTCATCTcgtacgctgcaggtcgac |
| med14-r-deg | TCTCCTAAGGGATAGTAGCGCCGGTGACATTTTATTCGCTatcgatgaattcgagctcg |
| pop2-f-deg | CAAGTACCAAGGTGTCATATACGGTATTGATGGGGACCAAcgtacgctgcaggtcgac |
| pop2-r-deg | TTTTTTTTTAAAATTGTGTATACATATAGTACATAAATGAcgatgaattcgagctcg |
| fcp1-f-deg | TTCGCAGTTGGAGGAAGAGTTGATGGATATGCTGGATGATcgtacgctgcaggtcgac |
| fcp1-r-deg | CAATGAGGAAAATGTGTGGAAAGATACGGCATCTGAGCTGcgatgaattcgagctcg |
| Pap2-AID-fwd | CGAAGATGATGATGAAGATGGATATAATCCTTATACCCTTCGTACGCTGCAGgtcgac |
| Pap2-AID-rev | ATGTACAGTTCAGTGCATCATTTAAACAAAAAGGCACATAATCGATGAATTCGAGCTCG |
| Nrd1-AID-fwd | GAATATGCTTAACCAACAGCAGCAGCAACAACAACAAAGCCGTACGCTGCAGgtcgac |
| Nrd1-AID-rev | TTTTATGTACTATGAGCAAATAAAGGGTGGAGTAAAGATCATCGATGAATTCGAGCTCG |
| Sen1-AID-fwd | ATCTAGCCCATTTATCCCAAAAAAAAGAAAGCCTAGATCACGTACGCTGCAGgtcga |
| Sen1-AID-rev | TATATATGCAGGTATAATTCCTAACACTTTTACTTCAAGAATCGATGAATTCGAGCTCG |
| Nab3-AID-fwd | TGTTCAAAGTCTATTAGATAGTTTAGCAAAACTACAAAAACGTACGCTGCAGgtcgac |
| Nab3-AID-rev | TATAATGTACAAGAAATGGAAAAGATTGAAAAAAGGGAGTATCGATGAATTCGAGCTCG |
| ccr4-f-deg | ATTTGAATTTATGAAGACAAACACAGGCAGTAAGAAAGTAcgtacgctgcaggtcgac |
| ccr4-r-deg | GTACAGAGAGGAGGGAGGGAGTGGATGAAAGTGTGCGGTatcgatgaattcgagctcg |
| not1_deg_f | CACCATCAATAGAAGGCAAACCCCTCTACAATCCAACGCAcgtacgctgcaggtcgac |
| not1_deg_r | CTGAAATCATGATTTTCGTATATAAATAAATGCAGTTTTTatcgatgaattcgagctcg |

### Table S3. FISH probes

Provided as an external excel spreadsheet.

### Supplementary Notes

#### Half-life estimation from 4tU labeling data

The following section describes the process by which we verified and tested various aspects of the half-life estimation procedures (Figures 1-2).

First we fitted binomial mixture models with a varying number of components, and found that 2 components were sufficient to describe the observed distribution of T→C conversions (Figure S1D). This model admits 3 parameters:

- The nascent fraction - $p_n$
- The background T→C conversion rate - ε
- The real T→C conversion rate ξ

Different samples along a 4tU labeling time course will obviously differ by their nascent fraction, but we wanted to verify that this is the only major difference between samples, i.e. that the error and conversion rate is common within an experimental batch and needs to be fitted once. We therefore fitted the data from the 4tU labeling time course allowing for each sample to have an individual set of parameters, or requiring samples to shae the error rate and the error+conversion rates (Figure S1E). In all cases, the nascent fraction was the single fitted parameter that varied the most, suggesting that the other parameters are much less time dependent. Furthermore, the constraint that samples will share the error and conversion rates caused only minor differences between the estimated nascent fraction, which is the critical parameter. This convinced us that these parameters (error and conversion rates) can be fitted globally once per experiment.

Next, when we fit the data with a 2-BMM model, we observe a linear increase in the fitted nascent fraction ($p_n$, Figure S1F), albeit with a time lag of ~1.5 minutes. Given that the nascent fraction should accumulate linearly for short labeling periods (assuming the model described in Methods section in main text), we interpret this result to mean that there is a lag between the addition of the 4tU to samples and the measurements of nascent molecules. This could be due to an actual delay in the time it takes 4tU to sufficiently accumulate in nuclei, or due to polyadenylation and maturation time of the first labeled nascent molecules. In any event, we use the time course data and fitted delay in subsequent experiments as a baseline assumption, adjusting the degradation rate equation from the main methods to include a time offset ($t_0$):

$$\delta = -\frac{1}{t-t_0}\ln(1 - p_n) - \gamma \quad (1)$$

Having verified that the cells are not distrubed from steady-state growth within several minutes of 4tU labeling (Figures S1G-H), we fit the half-life of individual transcripts using the 4tU time course data. We perform a maximum likelihood estimation, iterating between fitting the global parameters (dilution rate, labeling lag time, and conversion rate) and transcript-specific parameters (half-life and error rate, which in this case we allowed to be transcript-specific). When this process converges to the maximum, we calculate a confidence interval for each half-life estimation assuming a quadratic log likelihood around the maximum, or a linear log likelihood if the maximum was constrained (i.e. too stable/volatile). These estimates are the ones shown in Figure 1F.

These half-life estimates were obtained using 6 replicates in 4 time points, but we want to estimate the half-life from a single measurement. We calculated the estimated halflife from individual measurements (i.e. no time course data), by fitting the nascent fraction per transcript using the 2-BMM model, and using equation 1 (assuming a constant error rate for all transcripts).

We found that a single measurement after 6'-10' minutes of 4tU labeling results in good agreement with the full time course estimate (pearson $r$ 0.66, 0.85 for 6', 10' respectively, Figure S1I). As expected, the degree of agreement increases as more data is available per transcript (e.g. to pearson $r$ 0.81, 0.94 at median expression quantile and up, Figure S1I). This suggests that to the degree that a single measurement is worse than a full time course estimate, it can mostly be explained by decreased data availability, rather than the dynamic aspect of the time course.

Next, to verify that the differences between the single measurement estimates and time course estimates were not driven by the extra free parameter per transcript (error rate), we plot the transcript specific fitted error rate, and the ratio between the degradation rate fitted with the time course and a 10' single measurement (Figure S1J). We could not find correlation, suggesting that transcript-specific error rates were not necessary for a better fit. Importantly, 98.3% of fitted transcript error rates were below 0.011, compared to a conversion rate of ~0.11, making the differences between error rates negligible in virtually all cases (Figure S1J).

Finally, as the fitted half-lives depend on global parameters (growth rate, labeling time) that are not directly estimated from the data, we wanted to examine their effects on individual half-life fit. This amounts to probing equation 1 with respect to $\gamma$ and $t$. This analysis (Figure S1L) revealed that for most transcripts, a 2-fold error in growth rate estimate is insignificant (at least in wildtype conditions), as the active degradation is much faster than dilution. However, if the effective labeling time is different from the presumed labeling time (e.g. if there is an initial delay in 4tU incorporation, or a delay until molecules are observed due to polyadenylation) then the estimates of most transcripts will be offset by a constant.

### Xrn1 knockout effects

We tried to explain the log-fold changes to RNA level, degradation rates, and production rates observed between the wildtype and Xrn1 deletion strain at steady state using various feature sets and models.
We consider several basic features: ORF length, ORF GC content, 5' GC content, 3' GC content , and 3' UTR motif score(Cheng et al., 2017). On top of these ("base") we add additional features - translation efficiency(Cheng et al., 2019), and various measures derived from tRNA translation efficiency index(Sabi et al., 2017). On top of these, we used DRIMust(Leibovich et al., 2013) to look for significantly occurring motifs in transcripts that are extreme in either response (41 such sequences were found, mostly in ORFs and 3'UTRs), and considered their occurrences as additional features (termed "DM" in figure S2). Aside from these we also used a heuristic to select kmer-counts (1-7mers) as features by calculating correlations to data, sampling, and using a sparse linear model (LASSO) as a way to assess feature importance. This was repeated on random sets of kmers until convergence to a small set of informative features. Kmers were calculated separately for each ORF, 200 bp upstream of ORF (5UTR), and 200 bp downstream of ORF (3UTR). In figure S2H "FS" indicates all of these features were used simultaneously for prediction. Subsets of these features were provided to a LASSO (or a random forest) procedure trying to regress the given features against the target vector (logFC in degradation/production/rna levels for xrn1 KO relative to WT, or log levels of KO or WT). Results for various optimal models are shown in Figure S2G. Specific feature combination performance is shown for the differences between KO and WT in Figure S2H, and informative kmers are shown in S2I.

### First order model predictions when degradation is reduced but production is not

Assuming a first order model, the equation governing mRNA levels, R(t), following parameter changes is (neglecting the effects of growth):

$$R(t) = \frac{\pi}{\delta} + \left(\frac{\pi'}{\delta'} - \frac{\pi}{\delta}\right)\left(1 - e^{-t\delta'}\right)$$

Where the primed parameters (') are the new ones. Notably, at R(0) the value is the old steady-state and at t=inf the value is the new steady state. Notably, if the production rate doesn't change at t=0 then π simply scales the entire equation, i.e. if we consider the changes relative to t=0 we will remove any effects of the production rate from the signal. (red purple right hand side, below). Therefore, we consider the changes from t=0 by calculating the difference, i.e.:

$$dR(t) = R(t) - \frac{\pi}{\delta} = (\frac{\pi'}{\delta'} - \frac{\pi}{\delta})(1 - e^{-t\delta'})$$

Or in terms of the slope:

$$S = \frac{1}{t}dR(t) = \frac{1}{t}(\frac{\pi'}{\delta'} - \frac{\pi}{\delta})(1 - e^{-t\delta'})$$

Now if we assume production is constant, we have:

$$\log S = \log \pi + \log\frac{1}{t}(\frac{1}{\delta'} - \frac{1}{\delta})(1 - e^{-t\delta'})$$

For relatively small new degradation rates (δ'), the right hand side of the expression can be significantly simplified as:

$$(\frac{1}{\delta'} - \frac{1}{\delta}) \approx \frac{1}{\delta'}$$

And as:

$$(1 - e^{-t\delta'}) \approx \delta' t$$

Therefore:

$$\log S \approx \log \pi + \log(\frac{1}{t}\frac{1}{\delta'}\delta' t) = \log \pi$$

To illustrate this point, plotted below are the absolute, difference, and relative values of such responses when various parameters are changed (legend). They are grouped by their parameters. The case with the same production rate and initial degradation rate that changes to a variety of degradation rates is in green, the case with various production rates and a relatively short new half life is in purple, and the case with various production rates and a relatively long new half life is in red - where there’s the highest correlation of slate slope to the production rate.

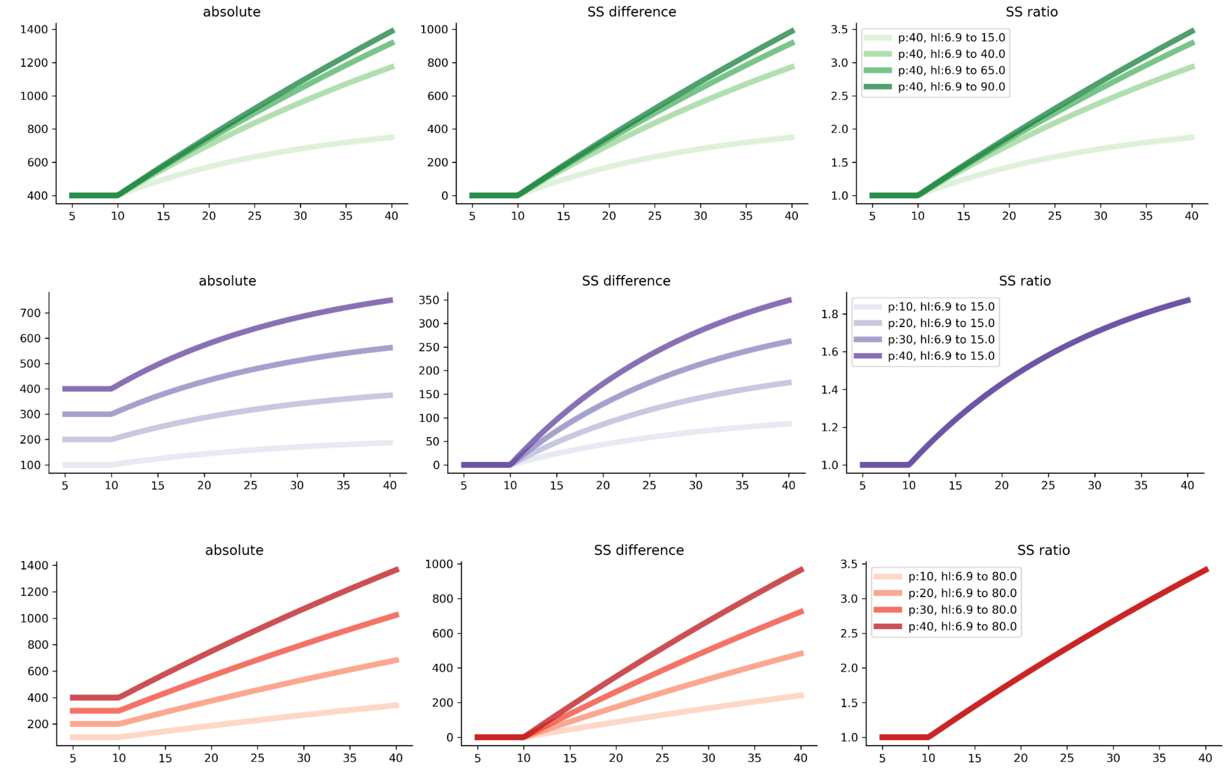


### Additional analyses

#### Not1 sensitive transcripts

mRNA trajectories were fitted with a pulse model, and the maximum upto t=120 was interpolated from the fitted model. Transcripts whose Not1 max value was larger than Xrn1 max value by at least a standard deviation of the wildtype max value (used as a null model) were considered “Not1 sensitive”.

### Supplementary References


Cheng, J., Maier, K.C., Avsec, Ž., Rus, P., and Gagneur, J. (2017). -regulatory elements explain most of the mRNA stability variation across genes in yeast. RNA *23*, 1648–1659.

Cheng, Z., Mugler, C.F., Keskin, A., Hodapp, S., Chan, L.Y.-L., Weis, K., Mertins, P., Regev, A., Jovanovic, M., and Brar, G.A. (2019). Small and Large Ribosomal Subunit Deficiencies Lead to Distinct Gene Expression Signatures that Reflect Cellular Growth Rate. Mol. Cell *73*, 36–47.e10.

Jonas, F., Soifer, I., and Barkai, N. (2018). A Visual Framework for Classifying Determinants of Cell Size. Cell Rep. *25*, 3519–3529.e2.

Jorgensen, P. (2002). Systematic Identification of Pathways That Couple Cell Growth and Division in Yeast. Science *297*, 395–400.

Leibovich, L., Paz, I., Yakhini, Z., and Mandel-Gutfreund, Y. (2013). DRIMust: a web server for discovering rank imbalanced motifs using suffix trees. Nucleic Acids Res. *41*, W174–W179.

Love, M.I., Huber, W., and Anders, S. (2014). Moderated estimation of fold change and dispersion for RNA-seq data with DESeq2. Genome Biology *15*.

Lu, A.X., Zarin, T., Hsu, I.S., and Moses, A.M. (2019). YeastSpotter: accurate and parameter-free web segmentation for microscopy images of yeast cells. Bioinformatics *35*, 4525–4527.

Sabi, R., Volvovitch Daniel, R., and Tuller, T. (2017). stAIcalc: tRNA adaptation index calculator based on species-specific weights. Bioinformatics *33*, 589–591.

Vallat, R. (2018). Pingouin: statistics in Python. Journal of Open Source Software *3*, 1026.

Zhang, J., Schneider, C., Ottmers, L., Rodriguez, R., Day, A., Markwardt, J., and Schneider, B.L. (2002). Genomic Scale Mutant Hunt Identifies Cell Size Homeostasis Genes in S. cerevisiae. Current Biology *12*, 1992–2001.
